# Multiple features within Sed5p mediate it’s COPI-independent trafficking and Golgi localization

**DOI:** 10.1101/765990

**Authors:** Guanbin Gao, David K. Banfield

## Abstract

Protein retention and the transport of proteins and lipids into and out of the Golgi is intimately linked to the biogenesis and homeostasis of this sorting hub of eukaryotic cells. Of particular importance are membrane proteins that mediate membrane fusion events with and within the Golgi – the Soluble N-ethylmaleimide-sensitive factor attachment protein receptors (SNAREs). In the Golgi of budding yeast cells a single syntaxin - the SNARE Sed5p - oversees membrane fusion within the Golgi. Determining how Sed5p is localized to and trafficked within the Golgi is critical to informing our understanding of the mechanism(s) of biogenesis and homeostasis of this organelle. Here we establish that the Golgi retention and trafficking of Sed5p between the Golgi and the ER is independent of COPI function, the composition of the transmembrane domain, and binding of the Sec1-Munc18 (SM) protein Sly1p. Rather, the steady state localization of Sed5p to the Golgi appears to be primarily conformation-based relying on intra-molecular associations between the Habc domain and SNARE-motif.

## INTRODUCTION

Within endomembrane network of membrane eukaryotic cells compartmental identity is generated through the interrelated processes of protein sorting and protein retention (Lee *et al*, 2004). Thus understanding how key protein constituents of endomembrane compartments are retained informs our understanding of organelle biogenesis and homeostasis - a matter of particular importance to understanding the membrane composition and biogenesis of the Golgi. The Golgi is the cell’s protein sorting hub: receiving and post-translationally modifying newly synthesized proteins; sorting proteins to topologically distal compartments or for retention within individual sub compartments (cisternae) of this organelle; recycling sorting machinery; and in returning errant ER resident proteins. In many respects the Golgi is unique among endomembrane organelles as it’s individual cisternae from *cis*, medial to *trans* harbor distinct enzyme compositions critical to the glycosylation of proteins and lipids – the glycosyltransferases and glycosidases. The mechanisms by which these enzymes are retained in the Golgi have been extensively studied (Banfield, 2011; Tu *et al*, 2008; Wood *et al*, 2009; Schmitz *et al*, 2008; Opat *et al*, 2001; Liu *et al*, 2018; Villeneuve *et al*, 2017). Moreover, following the distribution of Golgi membrane proteins by live cell imaging in yeast cells has provided critical insights into the biogenesis of this organelle (Losev *et al*, 2006; Matsuura-Tokita *et al*, 2006). The prevailing cisternal maturation model proposes that *cis* cisternae form *de novo* from ER-derived membranes, mature from *cis* to *trans*, then eventually disperse into vesicles (Glick *et al.*, 1997). Cisternal maturation is achieved by constant retrograde flow of components such as enzymes and transport factors from *trans* to *cis* cisternae to the early ones (Glick *et al.*, 1997) – a process mediated by the COPI coat complex. Thus enzymes and transport factors that undergo constant retrograde flow define the identity of Golgi cisterna.

It therefore follows that the composition of endomembrane compartments – such as the Golgi - are influenced to a large degree by membrane trafficking events which involve membrane fusion – and in particular the fusion of vesicular carriers (transport vesicles) with their target membranes. Among the various transport factors, soluble N-ethylmaleimide-sensitive factor attachment protein receptors (SNAREs) are of great importance. SNARE proteins are not only the minimum machinery required for membrane fusion, but also contribute to the specificity of the vesicle - target membrane fusion reaction (Söllner *et al.*, 1993; Rothman and Warren, 1994; Weber *et al.*, 1998). It is therefore of particular significance that SNARE proteins be correctly localized.

Through both physiological and structural studies an understanding of the multifaceted requirements of SNARE protein sorting and retention has been gained (McNew *et al*, 1997; Rayner & Pelham, 1997; Munro, 1995; Pryor *et al*, 2008). However, relatively little is known about how Golgi SNAREs, and in particular members of the syntaxin family of SNAREs are retained in this organelle (Banfield *et al*, 1994; Tochio *et al*, 2001). Given the complexity of membrane trafficking events within the Golgi and their critical importance in maintaining organelle identity as well as sorting events into and out of the Golgi, this represents a significant shortfall in our knowledge.

Sed5p is the only Golgi resident syntaxin SNARE in budding yeast (Hardwick and Pelham, 1992; Banfield *et al.*, 1994). Like other members of the syntaxin family, Sed5p is a type II membrane protein comprised of an N-terminal regulatory domain (Habc domain) followed by SNARE-motif and the C-terminal transmembrane domain (Dulubova *et al.*, 1999; Yamaguchi *et al.*, 2002). The very N-terminus of Sed5p (aa 1 – 10) contains a high affinity binding site for the Sec1/Munc18-like (SM) protein family member Sly1p (Søgaard *et al.*, 1994; Grabowski and Gallwitz, 1997; Bracher and Weissenhorn, 2002; Yamaguchi *et al.*, 2002). The Sly1p binding site is followed by a three-helix bundle termed the Habc domain (Yamaguchi *et al.*, 2002). The Habc domain has been shown to fold-back and associate with the SNARE-motif to form the closed conformation thereby preventing the SNARE-motif from engaging in SNARE complex formation (Demircioglu *et al.*, 2014). Sed5p is unique among the syntaxin family in that it can form several distinct SNARE complexes including one that mediates ER to Golgi transport (containing Sec22p, Bet1p and Bos1p) (Sacher *et al.*, 1997; Parlati *et al.*, 2000; Tsui *et al.*, 2001) and one that mediates intra-Golgi retrograde trafficking (containing Ykt6p, Gos1p and Sft1p) (Banfield *et al.*, 1995; McNew *et al.*, 1998; Tsui *et al.*, 2001; Parlati *et al.*, 2002; Graf *et al.*, 2005). Understanding the mechanism(s) by which Sed5p is localized to and within the Golgi will therefore provide important insight into the biogenesis and maintenance of this organelle.

A previous study determined that Sed5p’s Golgi-localization signal is only partially encoded by its transmembrane domain (Banfield *et al.*, 1994). Thus, additional signals in the cytosolic portion of Sed5p must also exist. Here we describe a comprehensive investigation into the mechanism(s) of Sed5p Golgi localization and trafficking between the Golgi and ER. We identify Sly1p-binding to Sed5p as being important for the robust export of Sed5p from the ER and in modulating the intra-Golgi distribution of Sed5p by masking the tribasic COPI-coatomer binding motif on Sed5p (^18^KKR^20^). Furthermore we establish that the Golgi retention and trafficking of Sed5p requires that the protein adopt a folded-back conformation in which the N-terminal Habc-containing region binds the SNARE-motif. Thus the primary mechanism governing Sed5p localization is conformation based rather than based on it’s COPI coatomer-binding motif or the composition or length of the transmembrane domain.

## RESULTS

### The localization of Sed5p to the Golgi is dependent in part on the amino acid sequence of it’s transmembrane domain

A previous study (Banfield *et al*, 1994) established that the Golgi localization of Sed5p / Syntaxin5 involved both features (composition and length) of the transmembrane domain (TMD) as well as the cytoplasmic portion of the protein. In the case of Sed5p it was shown that a Sed5p-Sso1p TMD chimera could support the growth of cells lacking endogenous Sed5p (Banfield *et al*, 1994). Although the localization of the protein was not examined in yeast cells, when the Sed5p-TMD chimeras were expressed in mammalian cells both the functional and non-functional Sed5p-Sso1p^TMD^ chimeras localized to the Golgi (Banfield *et al*, 1994). To address the relative contribution of the TMD to Sed5p’s function and localization we re-examined these findings. We generated Sed5p chimeras with the TMDs of Sso1p and Snc1p (Figure 1A.) Sso1p and Snc1p are part of the SNARE complex that mediates the fusion of exocytic vesicles with the plasma membrane (Gerst *et al*, 1992; Gurunathan *et al*, 2000; Aalto *et al*, 1993; Jäntti *et al*, 2002; Nakanishi *et al*, 2006) and as such are predominantly localized there (Gurunathan *et al*, 2000; Valdez-Taubas & Pelham, 2003). When expressed as the sole source of Sed5p, mNeon-Sed5p-Sso1p^TMD^ was found in puncta reminiscent of the Golgi and Sed5p-Sso1p^TMD^ could support the growth of yeast cells (Figure 1B and C). By contrast, mNeon-Sed5p-Snc1p^TMD^ was localized to the cell surface and to the vacuole, and Sed5p-Snc1p^TMD^ could not support the growth of cells lacking *SED5* (Figure 1B and C). These findings contrast with those on the localization of Nyv1p (Wen et al., 2006) where the Snc1p TMD was not sufficient to redirect a Nyv1p-Snc1p^TMD^ chimera (GNS) (Reggiori *et al*, 2000) from the limiting membrane of the vacuole to the cell surface unless Nyv1p’s canonical AP3-binding motif was disrupted (Wen *et al*, 2006). Given that the TMDs of Sso1p and Snc1p are similar in composition as well as length (Figure 1A) the cytoplasmic retention signal in Sed5p, in the context of non-canonical TMDs, would seem to have a rather subtle effect on the localization of the protein to the Golgi.

**Figure 1.**
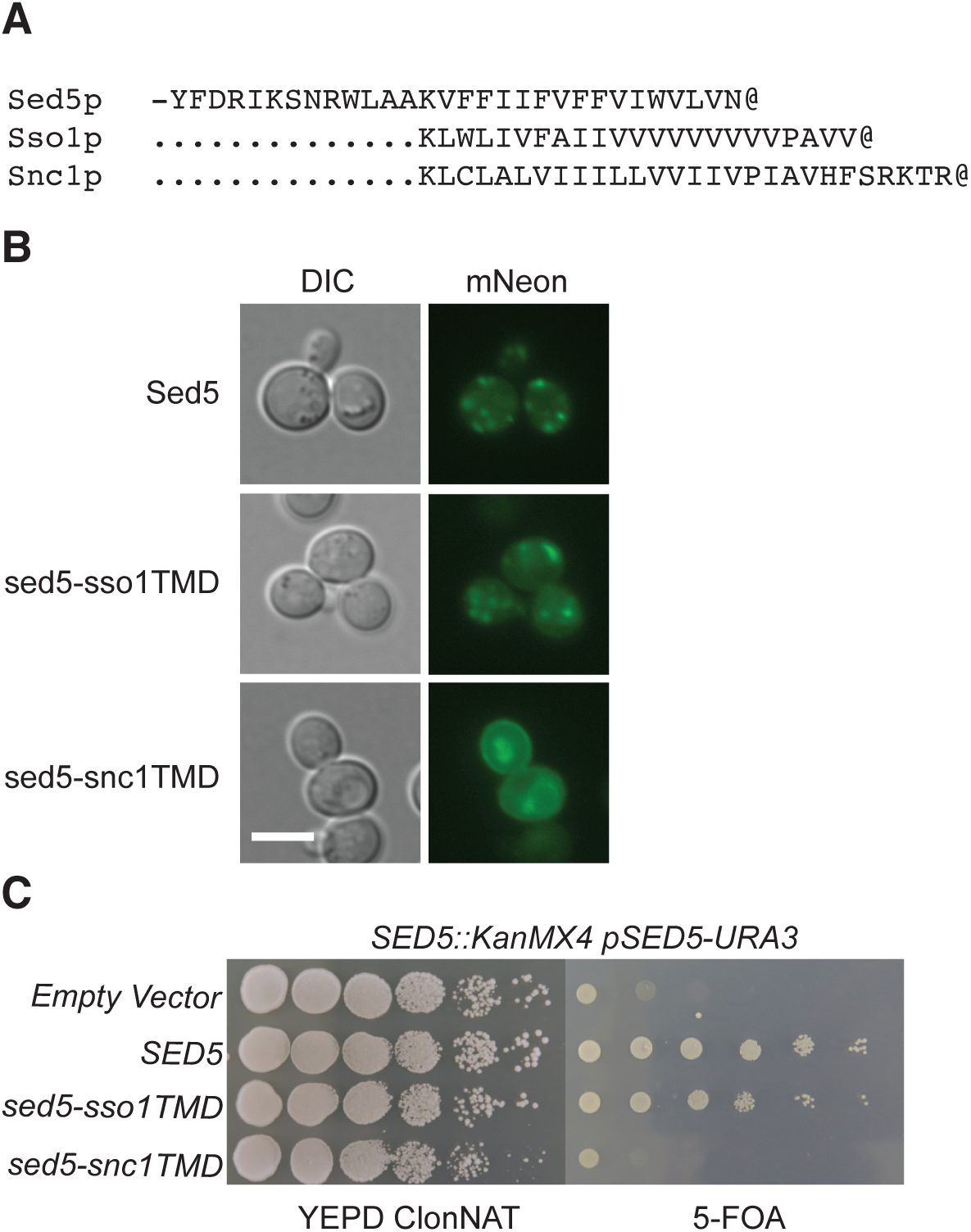
The transmembrane domain contributes to the Golgi retention of Sed5p. (A) Amino acid sequences of the transmembrane domains of Sed5p, Sso1p and Snc1p used to construct the Sed5-Sso1p and Sed5-Snc1p chimeras. The @ symbol denotes the C-terminus of the proteins (B) Fluorescence images from wild-type yeast cells expressing either wild-type mNeon-Sed5p or the mNeon-Sed5-Sso1p, mNeon-Sed5-Snc1p chimeras. Scale bar: 5 μm. (C) Functional assays of the *SED5-SSO1* and *SED5-SNC1* chimeras in yeast cells lacking endogenous *SED5*. Ten-fold serial dilutions (from left – right) of the indicated yeast transformants were spotted onto the surface of plates containing either YEPD-ClonNAT (for plasmid selection) or 5-FOA (to counter-select against the wild-type *SED5* gene, pSED5-URA3). Plates were incubated at 25°C for 2 days and then photographed.

### Sed5p possesses a tribasic COPI-binding motif in its N-terminus

Some yeast Golgi membrane proteins, including Golgi SNAREs, are known to cycle between the Golgi and the ER (Ballensiefen *et al*, 1998) and Sed5p has been shown to cycle between the Golgi and the ER constitutively (Wooding & Pelham, 1998). Moreover, Sed5p has also been shown to be incorporated into COPI coated vesicles in *in vitro* assays where a single substitution (S317A) significantly improved this incorporation - suggesting a role for phosphorylation in regulating the incorporation of Sed5p into COPI coated vesicles (Weinberger *et al*, 2005). However, identification of the features / sorting motifs on Sed5p responsible the protein’s retention and Golgi - ER cycling have remained elusive.

As a first step in the identification of signals on Sed5p that mediate COPI-based Golgi – ER transport we examined whether Sed5p could directly interact with COPI-coatomer. COPI-coatomer was purified by tandem affinity purification from cells expressing a Sec27p (β’-COP) fusion protein bearing a C-terminal calmodulin binding peptide (CBP) and a hexa-histidine sequence (see Table 1). Purified COPI-coatomer was mixed with bacterially expressed purified Sed5-GST and binding determined by immuno-staining proteins purified on GSH-beads with an anti-COPI-coatomer antibody (Figure 2A). Following a series of similar experiments, performed with truncated forms of wild-type and amino acid substituted Sed5p, we established that residues ^18^KKR^20^ accounted for the observed COPI-coatomer binding (Figure 2B). Amino acid substitutions at ^18^KKR^20^ (^18^AAA^20^), hereafter referred to a sed5p(KKR), abolished the interaction between Sed5p and COPI-coatomer *in vitro* whereas amino acid substitutions to the adjacent sequence ^21^NKNFR^25^ (^21^AAAAA^25^) had no effect (Figure 2B). The Sed5p COPI-coatomer binding “KKR” motif is highly conserved in yeasts (Figure 2C), and a similar motif is also present in Sed5p’s mammalian homolog Syntaxin5 (Hui *et al*, 1997). Syntaxin5 has two isoforms, the 35kDa short form (Syn5S) and 42kDa long form (Syn5L) (Figure 2C). The long form possesses an N-terminal extension relative to the short form that contains a tripeptide basic amino acid motif (^4^RKR^6^, Figure 2C). While the ^4^RKR^6^ motif in Stx5L has been shown to mediate the protein’s ER localization, the mechanism by which this occurs has not been explored (Hui *et al*, 1997). By contrast, the short form (Syn5S), which lacks the ^4^RKR^6^ motif, is primarily localized to the *cis* Golgi (Miyazaki *et al*, 2012). Given our findings with Sed5p we next asked whether the ^4^RKR^6^ motif in Syn5L could mediate binding to mammalian COPI-coatomer. As anticipated, bacterially expressed amino acids 1– 55 of Syn5L was able to bind to mammalian COPI-coatomer from whole cell homogenates (β-COP, Figure 1D), while this interaction was undetectable when ^4^RKR^6^ of Syn5L was substituted with alanine (^4^AAA^6^, Figure 1D). Thus, Syn5L’s steady-state localization to the ER is likely due to the protein’s retrieval from the Golgi (cis Golgi network) via its COPI-binding motif (^4^RKR^6^).

**Figure 2.**
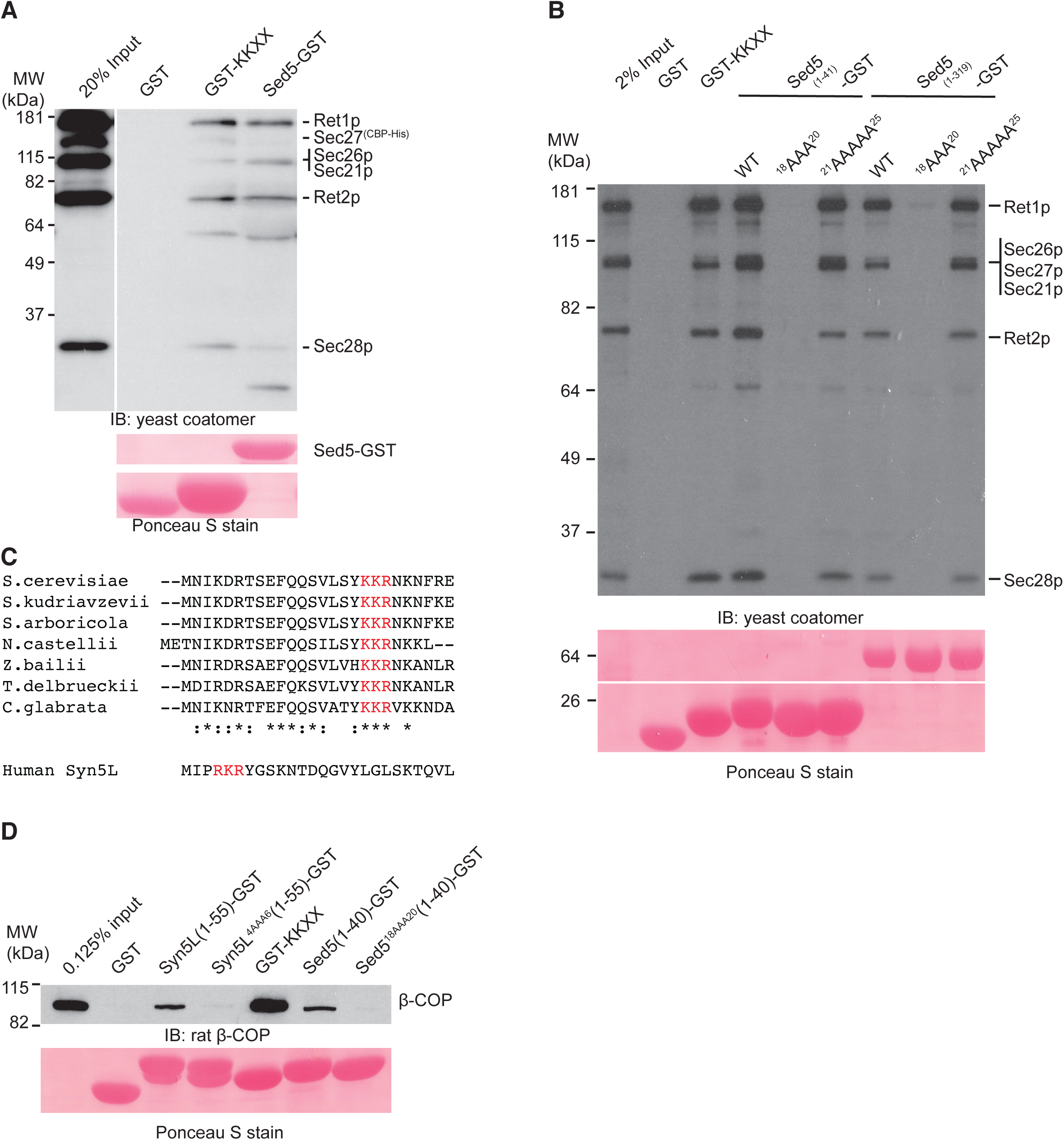
A tribasic motif in the N-terminus of Sed5p / Syntaxin 5L mediates binding to COPI-coatmer. (A) Sed5-GST binding to purified yeast COPI-coatomer. Purified bacterially expressed fusion proteins were mixed with purified yeast COPI-coatomer and bound proteins resolved by SDS-PAGE. GST and GST-KKXX serve as negative and positive controls, respectively. Following electrophoretic transfer to membranes purified recombinant proteins used in mixing assays were visualized with Ponceau S and thereafter membranes were immuno-stained (IB) with an anti-yeast COPI-coatomer antibody. (B) A tribasic motif “K18, K19, R20” in the N-terminus of Sed5p mediates binding to COPI-coatomer *in vitro*. The indicated GST-fusion proteins were mixed with yeast whole cell extracts and bound proteins resolved by SDS-PAGE followed by immuno-staining with an anti-yeast COPI-coatomer antibody (IB). GST and GST-KKXX serve as negative and positive controls, respectively. Purified recombinant proteins used in mixing assays were visualized with Ponceau S following electrophoretic transfer to membranes. (C) Amino acid sequences of the N-termini of fungal Sed5p proteins and the long form of human Syntaxin 5 (Syn5L). The tribasic motifs are indicated in red font. (D) The tribasic motif in Sed5p and the long form of human Syntaxin 5 mediate binding to human COPI-coatomer. The indicated purified GST-fusion proteins were mixed with whole cell extracts from 293T cells, and bound proteins were resolved by SDS-PAGE followed by immuno-staining an anti-βCOP antibody (IB). Following electrophoretic transfer to membranes purified recombinant proteins used in mixing assays were visualized with Ponceau S.

**Table 1.**
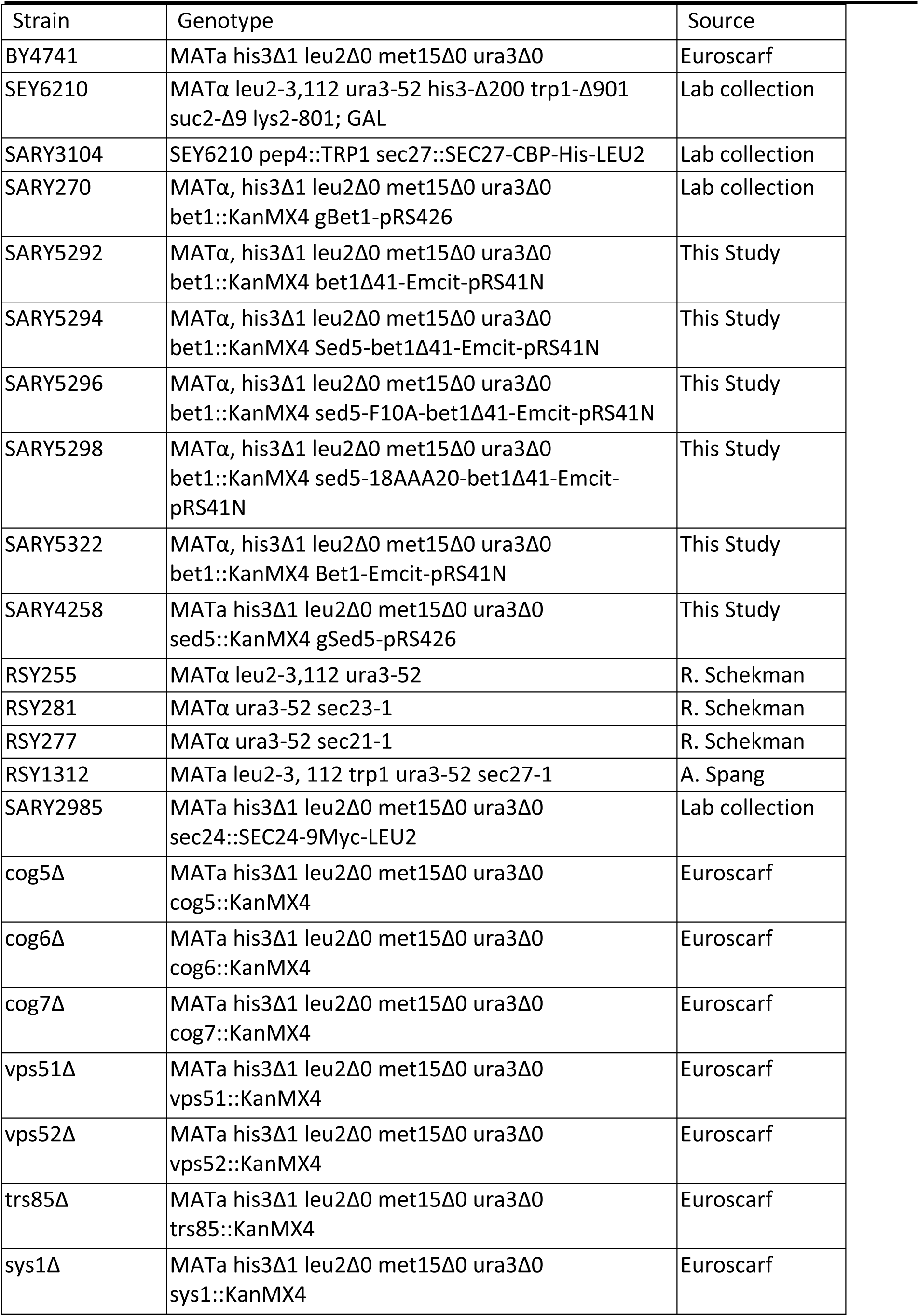
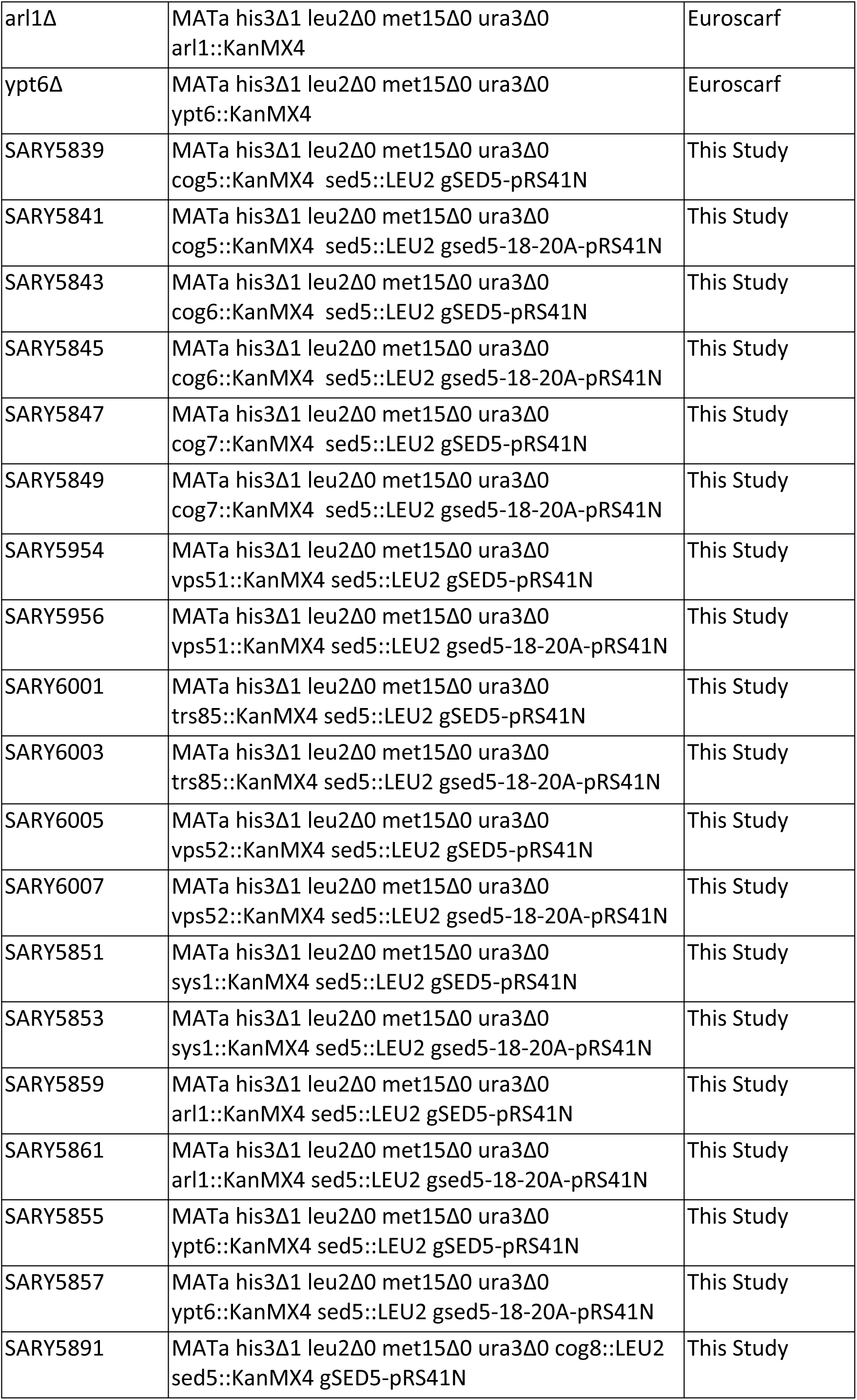
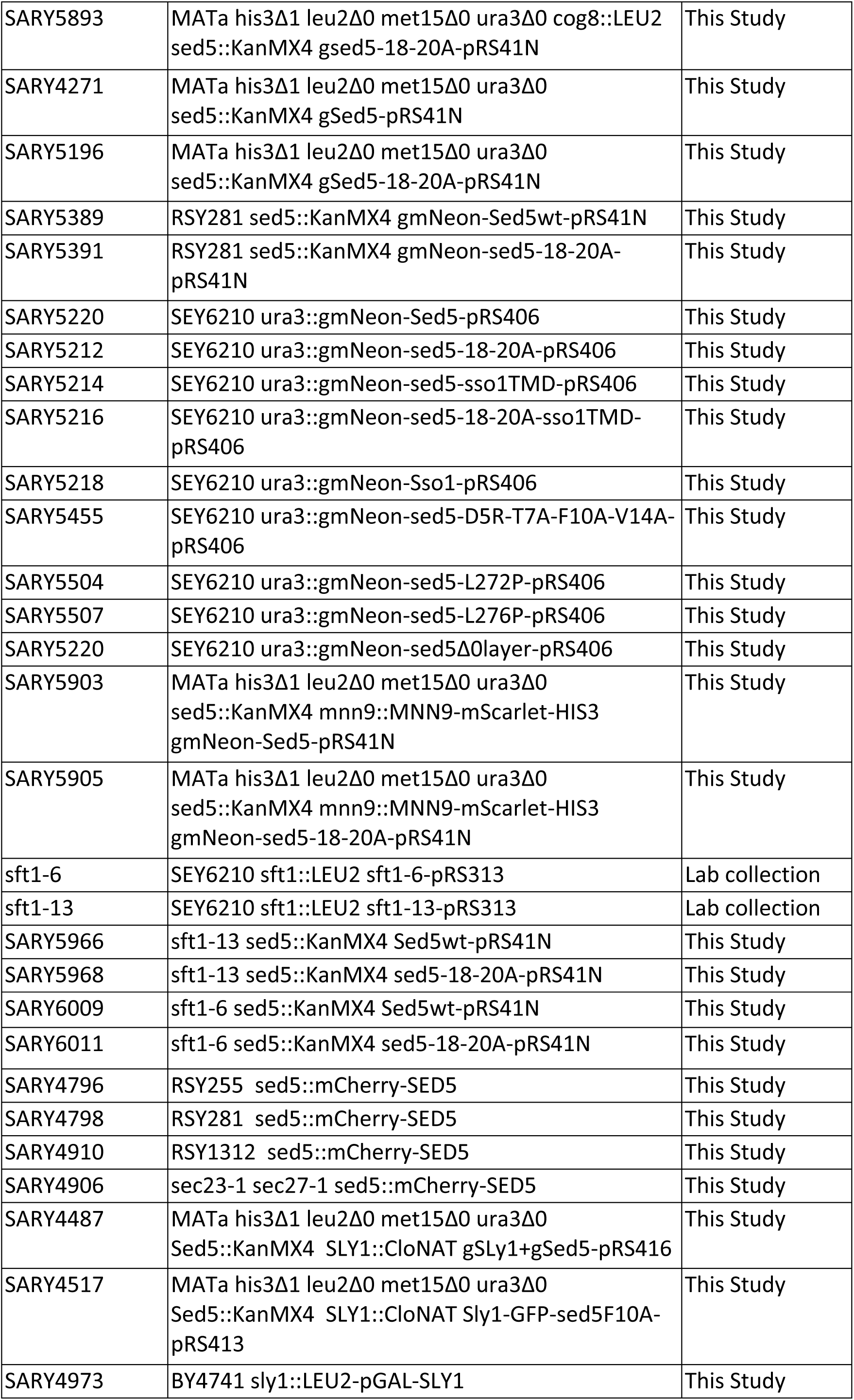
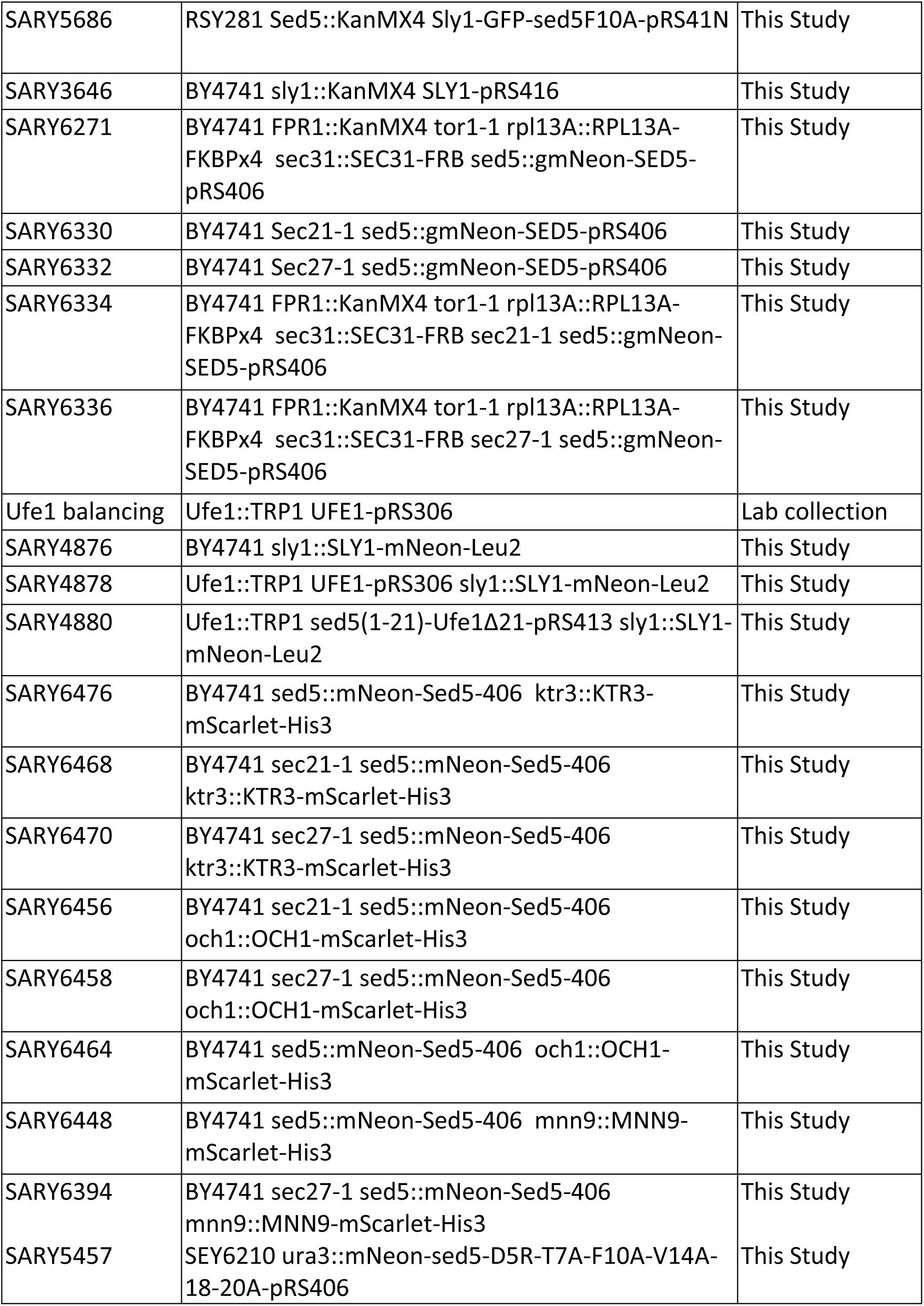
List of yeast strains used in this study.

| Strain | Genotype | Source |
| --- | --- | --- |
| BY4741 | MATa his3Δ1 leu2Δ0 met15Δ0 ura3Δ0 | Euroscarf |
| SEY6210 | MATα leu2-3,112 ura3-52 his3-Δ200 trp1-Δ901<br>suc2-Δ9 lys2-801; GAL | Lab collection |
| SARY3104 | SEY6210 pep4::TRP1 sec27::SEC27-CBP-His-LEU2 | Lab collection |
| SARY270 | MATα, his3Δ1 leu2Δ0 met15Δ0 ura3Δ0<br>bet1::KanMX4 gBet1-pRS426 | Lab collection |
| SARY5292 | MATα, his3Δ1 leu2Δ0 met15Δ0 ura3Δ0<br>bet1::KanMX4 bet1Δ41-Emcit-pRS41N | This Study |
| SARY5294 | MATα, his3Δ1 leu2Δ0 met15Δ0 ura3Δ0<br>bet1::KanMX4 Sed5-bet1Δ41-Emcit-pRS41N | This Study |
| SARY5296 | MATα, his3Δ1 leu2Δ0 met15Δ0 ura3Δ0<br>bet1::KanMX4 sed5-F10A-bet1Δ41-Emcit-pRS41N | This Study |
| SARY5298 | MATα, his3Δ1 leu2Δ0 met15Δ0 ura3Δ0<br>bet1::KanMX4 sed5-18AAA20-bet1Δ41-Emcit-pRS41N | This Study |
| SARY5322 | MATα, his3Δ1 leu2Δ0 met15Δ0 ura3Δ0<br>bet1::KanMX4 Bet1-Emcit-pRS41N | This Study |
| SARY4258 | MATa his3Δ1 leu2Δ0 met15Δ0 ura3Δ0<br>sed5::KanMX4 gSed5-pRS426 | This Study |
| RSY255 | MATα leu2-3,112 ura3-52 | R. Schekman |
| RSY281 | MATα ura3-52 sec23-1 | R. Schekman |
| RSY277 | MATα ura3-52 sec21-1 | R. Schekman |
| RSY1312 | MATa leu2-3, 112 trp1 ura3-52 sec27-1 | A. Spang |
| SARY2985 | MATa his3Δ1 leu2Δ0 met15Δ0 ura3Δ0<br>sec24::SEC24-9Myc-LEU2 | Lab collection |
| cog5Δ | MATa his3Δ1 leu2Δ0 met15Δ0 ura3Δ0<br>cog5::KanMX4 | Euroscarf |
| cog6Δ | MATa his3Δ1 leu2Δ0 met15Δ0 ura3Δ0<br>cog6::KanMX4 | Euroscarf |
| cog7Δ | MATa his3Δ1 leu2Δ0 met15Δ0 ura3Δ0<br>cog7::KanMX4 | Euroscarf |
| vps51Δ | MATa his3Δ1 leu2Δ0 met15Δ0 ura3Δ0<br>vps51::KanMX4 | Euroscarf |
| vps52Δ | MATa his3Δ1 leu2Δ0 met15Δ0 ura3Δ0<br>vps52::KanMX4 | Euroscarf |
| trs85Δ | MATa his3Δ1 leu2Δ0 met15Δ0 ura3Δ0<br>trs85::KanMX4 | Euroscarf |
| sys1Δ | MATa his3Δ1 leu2Δ0 met15Δ0 ura3Δ0<br>sys1::KanMX4 | Euroscarf |
| arl1Δ | MATa his3Δ1 leu2Δ0 met15Δ0 ura3Δ0<br>arl1::KanMX4 | Euroscarf |
| ypt6Δ | MATa his3Δ1 leu2Δ0 met15Δ0 ura3Δ0<br>ypt6::KanMX4 | Euroscarf |
| SARY5839 | MATa his3Δ1 leu2Δ0 met15Δ0 ura3Δ0<br>cog5::KanMX4 sed5::LEU2 gSED5-pRS41N | This Study |
| SARY5841 | MATa his3Δ1 leu2Δ0 met15Δ0 ura3Δ0<br>cog5::KanMX4 sed5::LEU2 gsed5-18-20A-pRS41N | This Study |
| SARY5843 | MATa his3Δ1 leu2Δ0 met15Δ0 ura3Δ0<br>cog6::KanMX4 sed5::LEU2 gSED5-pRS41N | This Study |
| SARY5845 | MATa his3Δ1 leu2Δ0 met15Δ0 ura3Δ0<br>cog6::KanMX4 sed5::LEU2 gsed5-18-20A-pRS41N | This Study |
| SARY5847 | MATa his3Δ1 leu2Δ0 met15Δ0 ura3Δ0<br>cog7::KanMX4 sed5::LEU2 gSED5-pRS41N | This Study |
| SARY5849 | MATa his3Δ1 leu2Δ0 met15Δ0 ura3Δ0<br>cog7::KanMX4 sed5::LEU2 gsed5-18-20A-pRS41N | This Study |
| SARY5954 | MATa his3Δ1 leu2Δ0 met15Δ0 ura3Δ0<br>vps51::KanMX4 sed5::LEU2 gSED5-pRS41N | This Study |
| SARY5956 | MATa his3Δ1 leu2Δ0 met15Δ0 ura3Δ0<br>vps51::KanMX4 sed5::LEU2 gsed5-18-20A-pRS41N | This Study |
| SARY6001 | MATa his3Δ1 leu2Δ0 met15Δ0 ura3Δ0<br>trs85::KanMX4 sed5::LEU2 gSED5-pRS41N | This Study |
| SARY6003 | MATa his3Δ1 leu2Δ0 met15Δ0 ura3Δ0<br>trs85::KanMX4 sed5::LEU2 gsed5-18-20A-pRS41N | This Study |
| SARY6005 | MATa his3Δ1 leu2Δ0 met15Δ0 ura3Δ0<br>vps52::KanMX4 sed5::LEU2 gSED5-pRS41N | This Study |
| SARY6007 | MATa his3Δ1 leu2Δ0 met15Δ0 ura3Δ0<br>vps52::KanMX4 sed5::LEU2 gsed5-18-20A-pRS41N | This Study |
| SARY5851 | MATa his3Δ1 leu2Δ0 met15Δ0 ura3Δ0<br>sys1::KanMX4 sed5::LEU2 gSED5-pRS41N | This Study |
| SARY5853 | MATa his3Δ1 leu2Δ0 met15Δ0 ura3Δ0<br>sys1::KanMX4 sed5::LEU2 gsed5-18-20A-pRS41N | This Study |
| SARY5859 | MATa his3Δ1 leu2Δ0 met15Δ0 ura3Δ0<br>arl1::KanMX4 sed5::LEU2 gSED5-pRS41N | This Study |
| SARY5861 | MATa his3Δ1 leu2Δ0 met15Δ0 ura3Δ0<br>arl1::KanMX4 sed5::LEU2 gsed5-18-20A-pRS41N | This Study |
| SARY5855 | MATa his3Δ1 leu2Δ0 met15Δ0 ura3Δ0<br>ypt6::KanMX4 sed5::LEU2 gSED5-pRS41N | This Study |
| SARY5857 | MATa his3Δ1 leu2Δ0 met15Δ0 ura3Δ0<br>ypt6::KanMX4 sed5::LEU2 gsed5-18-20A-pRS41N | This Study |
| SARY5891 | MATa his3Δ1 leu2Δ0 met15Δ0 ura3Δ0 cog8::LEU2<br>sed5::KanMX4 gSED5-pRS41N | This Study |
| SARY5893 | MATa his3Δ1 leu2Δ0 met15Δ0 ura3Δ0 cog8::LEU2 sed5::KanMX4 gsed5-18-20A-pRS41N | This Study |
| SARY4271 | MATa his3Δ1 leu2Δ0 met15Δ0 ura3Δ0 sed5::KanMX4 gSed5-pRS41N | This Study |
| SARY5196 | MATa his3Δ1 leu2Δ0 met15Δ0 ura3Δ0 sed5::KanMX4 gSed5-18-20A-pRS41N | This Study |
| SARY5389 | RSY281 sed5::KanMX4 gmNeon-Sed5wt-pRS41N | This Study |
| SARY5391 | RSY281 sed5::KanMX4 gmNeon-sed5-18-20A-pRS41N | This Study |
| SARY5220 | SEY6210 ura3::gmNeon-Sed5-pRS406 | This Study |
| SARY5212 | SEY6210 ura3::gmNeon-sed5-18-20A-pRS406 | This Study |
| SARY5214 | SEY6210 ura3::gmNeon-sed5-sso1TMD-pRS406 | This Study |
| SARY5216 | SEY6210 ura3::gmNeon-sed5-18-20A-sso1TMD-pRS406 | This Study |
| SARY5218 | SEY6210 ura3::gmNeon-Sso1-pRS406 | This Study |
| SARY5455 | SEY6210 ura3::gmNeon-sed5-D5R-T7A-F10A-V14A-pRS406 | This Study |
| SARY5504 | SEY6210 ura3::gmNeon-sed5-L272P-pRS406 | This Study |
| SARY5507 | SEY6210 ura3::gmNeon-sed5-L276P-pRS406 | This Study |
| SARY5220 | SEY6210 ura3::gmNeon-sed5Δ0layer-pRS406 | This Study |
| SARY5903 | MATa his3Δ1 leu2Δ0 met15Δ0 ura3Δ0 sed5::KanMX4 mnn9::MNN9-mScarlet-HIS3 gmNeon-Sed5-pRS41N | This Study |
| SARY5905 | MATa his3Δ1 leu2Δ0 met15Δ0 ura3Δ0 sed5::KanMX4 mnn9::MNN9-mScarlet-HIS3 gmNeon-sed5-18-20A-pRS41N | This Study |
| sft1-6 | SEY6210 sft1::LEU2 sft1-6-pRS313 | Lab collection |
| sft1-13 | SEY6210 sft1::LEU2 sft1-13-pRS313 | Lab collection |
| SARY5966 | sft1-13 sed5::KanMX4 Sed5wt-pRS41N | This Study |
| SARY5968 | sft1-13 sed5::KanMX4 sed5-18-20A-pRS41N | This Study |
| SARY6009 | sft1-6 sed5::KanMX4 Sed5wt-pRS41N | This Study |
| SARY6011 | sft1-6 sed5::KanMX4 sed5-18-20A-pRS41N | This Study |
| SARY4796 | RSY255 sed5::mCherry-SED5 | This Study |
| SARY4798 | RSY281 sed5::mCherry-SED5 | This Study |
| SARY4910 | RSY1312 sed5::mCherry-SED5 | This Study |
| SARY4906 | sec23-1 sec27-1 sed5::mCherry-SED5 | This Study |
| SARY4487 | MATa his3Δ1 leu2Δ0 met15Δ0 ura3Δ0 Sed5::KanMX4 SLY1::CloNAT gSLy1+gSed5-pRS416 | This Study |
| SARY4517 | MATa his3Δ1 leu2Δ0 met15Δ0 ura3Δ0 Sed5::KanMX4 SLY1::CloNAT Sly1-GFP-sed5F10A-pRS413 | This Study |
| SARY4973 | BY4741 sly1::LEU2-pGAL-SLY1 | This Study |
| SARY5686 | RSY281 Sed5::KanMX4 Sly1-GFP-sed5F10A-pRS41N | This Study |
| SARY3646 | BY4741 sly1::KanMX4 SLY1-pRS416 | This Study |
| SARY6271 | BY4741 FPR1::KanMX4 tor1-1 rpl13A::RPL13A-FKBPx4 sec31::SEC31-FRB sed5::gmNeon-SED5-pRS406 | This Study |
| SARY6330 | BY4741 Sec21-1 sed5::gmNeon-SED5-pRS406 | This Study |
| SARY6332 | BY4741 Sec27-1 sed5::gmNeon-SED5-pRS406 | This Study |
| SARY6334 | BY4741 FPR1::KanMX4 tor1-1 rpl13A::RPL13A-FKBPx4 sec31::SEC31-FRB sec21-1 sed5::gmNeon-SED5-pRS406 | This Study |
| SARY6336 | BY4741 FPR1::KanMX4 tor1-1 rpl13A::RPL13A-FKBPx4 sec31::SEC31-FRB sec27-1 sed5::gmNeon-SED5-pRS406 | This Study |
| Ufe1 balancing | Ufe1::TRP1 UFE1-pRS306 | Lab collection |
| SARY4876 | BY4741 sly1::SLY1-mNeon-Leu2 | This Study |
| SARY4878 | Ufe1::TRP1 UFE1-pRS306 sly1::SLY1-mNeon-Leu2 | This Study |
| SARY4880 | Ufe1::TRP1 sed5(1-21)-Ufe1Δ21-pRS413 sly1::SLY1-mNeon-Leu2 | This Study |
| SARY6476 | BY4741 sed5::mNeon-Sed5-406 ktr3::KTR3-mScarlet-His3 | This Study |
| SARY6468 | BY4741 sec21-1 sed5::mNeon-Sed5-406 ktr3::KTR3-mScarlet-His3 | This Study |
| SARY6470 | BY4741 sec27-1 sed5::mNeon-Sed5-406 ktr3::KTR3-mScarlet-His3 | This Study |
| SARY6456 | BY4741 sec21-1 sed5::mNeon-Sed5-406 och1::OCH1-mScarlet-His3 | This Study |
| SARY6458 | BY4741 sec27-1 sed5::mNeon-Sed5-406 och1::OCH1-mScarlet-His3 | This Study |
| SARY6464 | BY4741 sed5::mNeon-Sed5-406 och1::OCH1-mScarlet-His3 | This Study |
| SARY6448 | BY4741 sed5::mNeon-Sed5-406 mnn9::MNN9-mScarlet-His3 | This Study |
| SARY6394 | BY4741 sec27-1 sed5::mNeon-Sed5-406 mnn9::MNN9-mScarlet-His3 | This Study |
| SARY5457 | SEY6210 ura3::mNeon-sed5-D5R-T7A-F10A-V14A-18-20A-pRS406 | This Study |

**Table 2.**
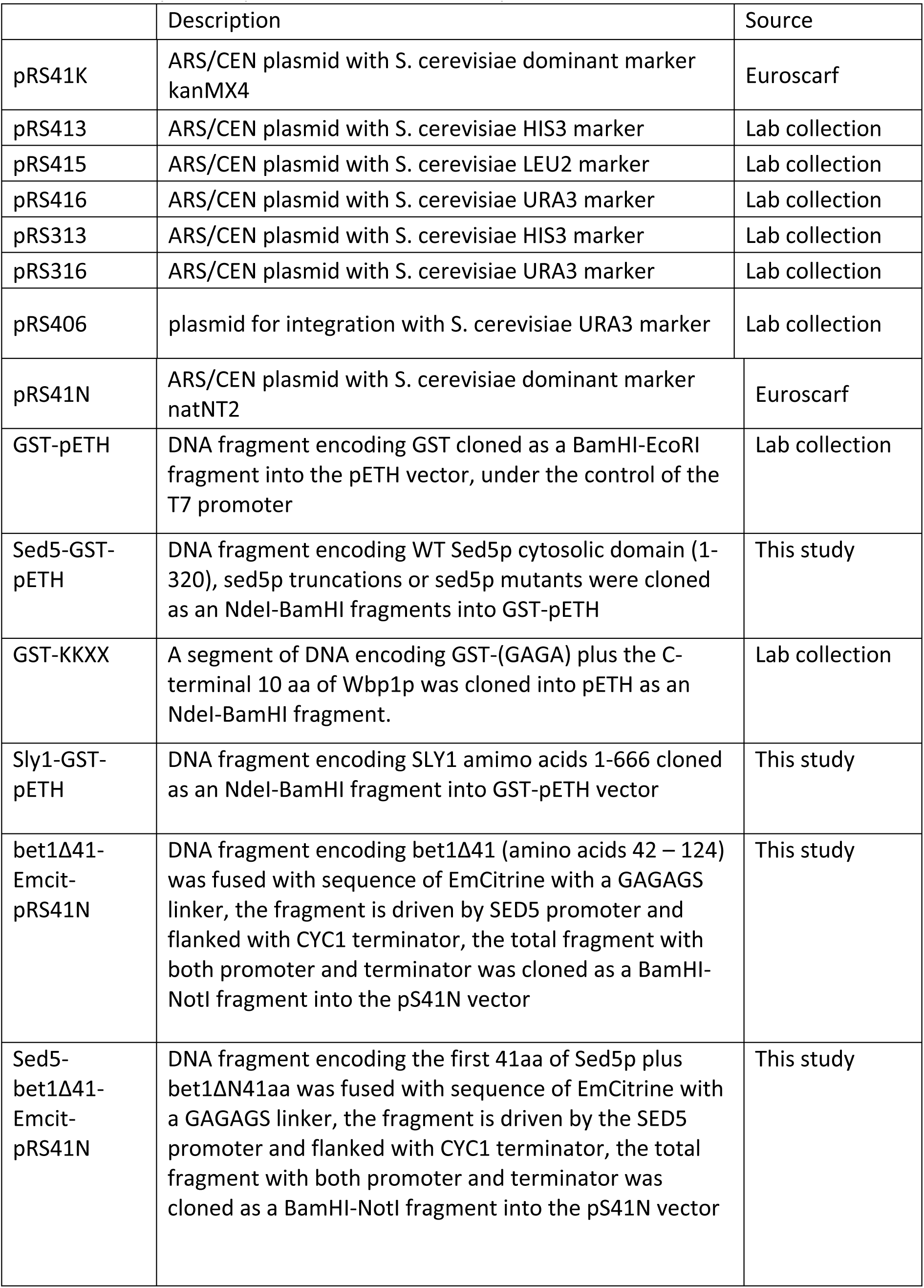
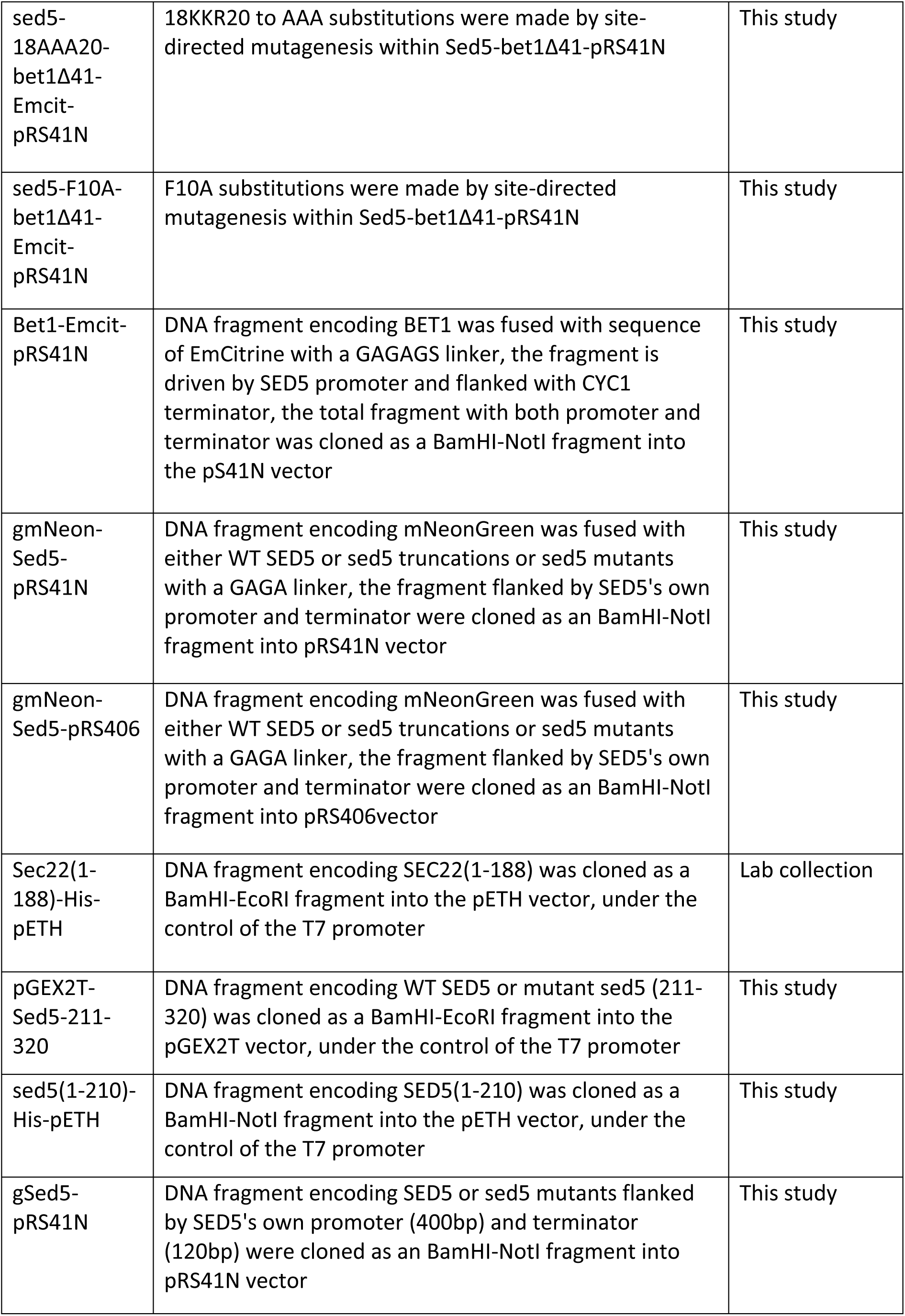
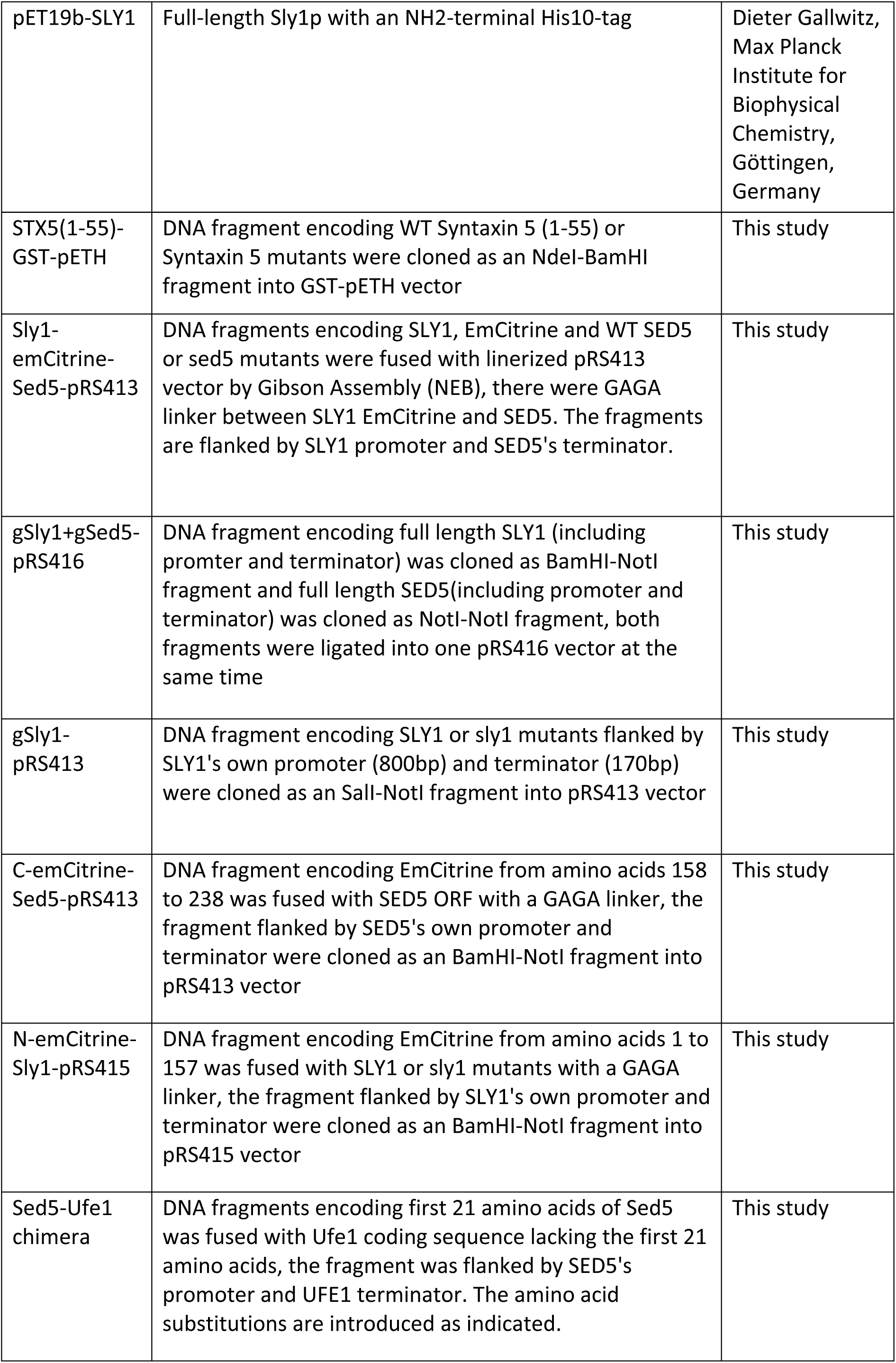
Description of plasmids used in this study.

To date relatively few N-terminal COPI-binding motifs have been characterized (Arakel & Schwappach, 2018; Tu *et al*, 2008; Liu *et al*, 2018). In mammalian cells, some Golgi resident glycosyltransferases have been shown to directly interact with coatomer (Liu *et al*, 2018) however the motif that mediates this association – “ϕ-(K/R)-X-L-X-(K/R)” - is not similar to the motif we have identified in Sed5p (Figure 2). The COPI-coatomer binding motif from Sed5p is also able to mediate binding to mammalian coatomer, and thus a binding site for N-terminal tri-basic motifs on COPI-coatomer would seem to be an evolutionarily conserved feature.

### The COPI coatomer-binding tribasic motif (KKR) is not required for the Golgi-ER trafficking of Sed5p

In yeast cells amino acid substitutions that ablate COPI coatomer-binding sites typically result in mislocalization of the altered membrane proteins to the vacuole (Sato *et al*, 2001). Having shown that the ^18^KKR^20^ motif of Sed5p was required for COPI coatomer-binding *in vitro*, we next investigated whether this motif was essential for the function of *SED5* and whether the ^18^KKR^20^ motif contributed to Sed5p’s Golgi localization. Cells expressing *sed5*(^18^KKR^20^) as their sole source of *SED5* are viable – a result that suggested that sed5p(^18^KKR^20^) was unlikely to be extensively mislocalized in cells (Figure 3A). Indeed, mNeon-sed5p(KKR) localized to puncta (Figure 3B). To determine the extent to which the localization of mNeon-sed5p(KKR) was altered in cells we used the *cis* Golgi resident glycosyltransferase Mnn9p as a co-localization marker. Wild-type mNeon-Sed5p showed ∼ 85% overlapping distribution with Mnn9p-mScarlett (Figure 3B and C) whereas mNeon-sed5p(KKR) showed a modest but statistically significant reduction in co-localization with Mnn9-mScarlett (∼70%). We saw no evidence of vacuolar localization (Figure 3B). Thus, the COPI coatomer-binding motif does not appear to be required for Sed5p’s Golgi retention *per se*.

**Figure 3.**
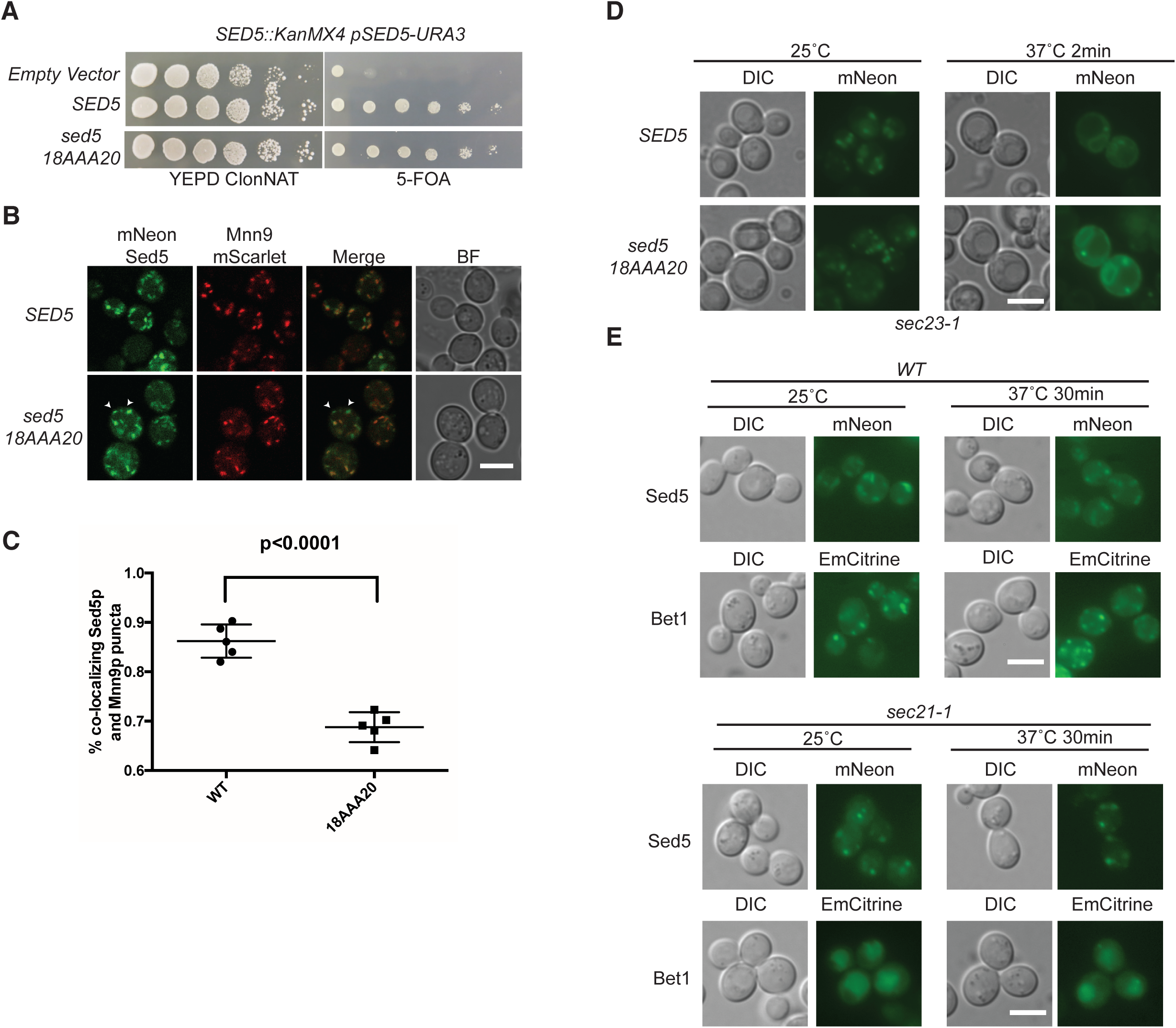
The N-terminal tribasic motif “K18,K19,R20” (KKR) mediates the intra-Golgi distribution of Sed5p. (A) *sed5* (KKR) can support the growth of cells lacking the wild-type *SED5* gene. (B) mNeon-sed5p(KKR) shows a reduced overlapping distribution with the *cis* Golgi marker Mnn9p-mScarlet. Wild-type yeast cells expressing a chromosomally tagged form of Mnn9p-mScarlet were transformed with either wild-type or amino acid substituted mNeon-sed5p as the only source of Sed5p. White arrowheads indicate puncta that do not coincide with Mnn9p-mScarlet. Scale bar 5 μm. (C) Transformants from the experiment depicted in (A) were assessed for co-localization between mNeon-Sed5p, mNeon-sed5p(KKR) and Mnn9p-mScarlet. The percentage of co-localizing puncta is plotted for mNeon-Sed5p (WT) and the tribasic amino acid substituted mNeon-sed5p(KKR). Mean ± SD *P < 0.0001 calculated by unpaired Student’s t-test assuming Unequal Variances; N = 5 isolates. n>1,000 cells. (D) Amino acid substituted Sed5p (mNeon-sed5p(KKR)) is not defective in Golgi – ER trafficking. The indicated genes were introduced into *sec23-1* cells as the only source of Sed5p, and transformants were grown at 25°C or 37°C (for 2 minutes) prior to being photographed. (E) mNeon-Sed5p is not mislocalized to the vacuole in a COPI-coatomer mutant (*sec21-1*) whereas the Qc-SNARE Bet1p-emCitrine is. The indicated genes were introduced into wild-type or *sec21-1* cells and transformants incubated at 25°C or 37°C prior to being photographed. Scale bar = 5 μm.

To determine whether the recycling of the COPI coatomer-binding deficient form of Sed5p (sed5p(KKR)) was deficient in retrograde transport from the Golgi to the ER we examined the fate of the protein in cells carrying the *sec23-1* mutation. When grown at 37°C *sec23-1* cells are defective in COPII vesicle formation and consequently protein export from the ER is blocked. However, a temperature-sensitive block in ER export however does not immediately impact Golgi-ER retrograde transport (Wooding & Pelham, 1998). Therefore if the COPI-binding motif in Sed5p (aa 18-20) accounted for the robust recycling of the protein (Wooding & Pelham, 1998) then amino acid substitutions to this motif would be expected to prevent the accumulation of sed5p(KKR) in the ER in *sec23-1* cells grown at 37°C. However, following a brief incubation at 37°C (2 minutes) the localization pattern of wild-type and amino acid substituted Sed5p (sed5p(KKR)) were comparable –although some Golgi-like puncta were still apparent in the case of mNeon-sed5p(KKR) (Figure 3D). Thus the COPI coatomer-binding motif in Sed5p does not substantially contribute to the robust redistribution of wild-type Sed5p from the Golgi to the ER observed in *sec23-1* cells. In agreement with these data we find that Sed5p was not mislocalized to the vacuole in COPI mutants (Figure 3E) unlike its cognate Qc-SNARE partner Bet1p (Figure 3E), and the Golgi-resident glycosyltransferases Ktr3p, Och1p and Mnn9p (Figure S1).

### Sed5p’s Golgi-ER trafficking is independent of COPI function

The data presented thus far indicated that Sed5p is rapidly retrieved to the ER in *sec23-1* cells at 37°C (Figure 3D), and yet Sed5p is not mislocalized from the Golgi in coatomer mutants or when its “KKR” COPI-coatomer binding motif is disrupted (Figure 3D and E). To seek additional insight into the mechanism by which Sed5p is retrieved to the ER we examined the fate of mCherry-Sed5 in cells carrying both COPI and COPII conditional mutations (Figure 4). Following a brief incubation of *sec23-1* cells at 37°C mCherry-Sed5p colocalized to the ER, whereas mCherry-Sed5p remained in puncta in *sec27-1* cells incubated at 37°C (Figure 4A). When mCherry-Sed5p was expressed in cells carrying both the *sec23-1* and *sec27-1* alleles the protein localized to puncta at 25°C and to the ER at 37°C (Figure 4A). These data are seemingly inconsistent with a role for COPI in the retrieval of Sed5p to the ER, however, if the inactivation of *sec23* proceeded more rapidly than that of *sec27* this might indirectly result in a block in retrograde transport by inhibiting ER – Golgi trafficking (Wooding & Pelham, 1998). To address this potential pitfall we modified the experimental design to incorporate a knock-sideways strategy (Papanikou *et al*, 2015). We generated yeast strains carrying either the *sec21-1* (γ-COP) allele or *sec27-1* (β’COP) allele in which the ribosomal protein Rpl13A was tagged with 4 copies of FK506 binding protein (FKBP) and *SEC31* was tagged with one copy of the fragment of mammalian target of rapamycin (mTOR) that binds FKBP12 (FRB) (see Table 1). Previous studies have shown that in this genetic background addition of rapamycin will cause Sec31p-FRB to be sequestered by Rpl13A-(FKBP)_4_ and hence COPII-mediated ER export will become blocked (Papanikou *et al*, 2015). In the absence of rapamycin mNeon-Sed5p was found in the Golgi at 25°C (Figure 4B). When cells were incubated at 37°C for 15 minutes (to inactivate COPI) prior to the addition of rapamycin or for 30 minutes prior to the addition of rapamycin (to inactivate COPII), mNeon-Sed5 was still localized to the ER after 15 minutes of rapamycin treatment (Figure 4B). Based on these, and other data presented thus far, we conclude that Sed5p’s Golgi-ER retrieval does not directly involve COPI-coatomer function.

**Figure 4.**
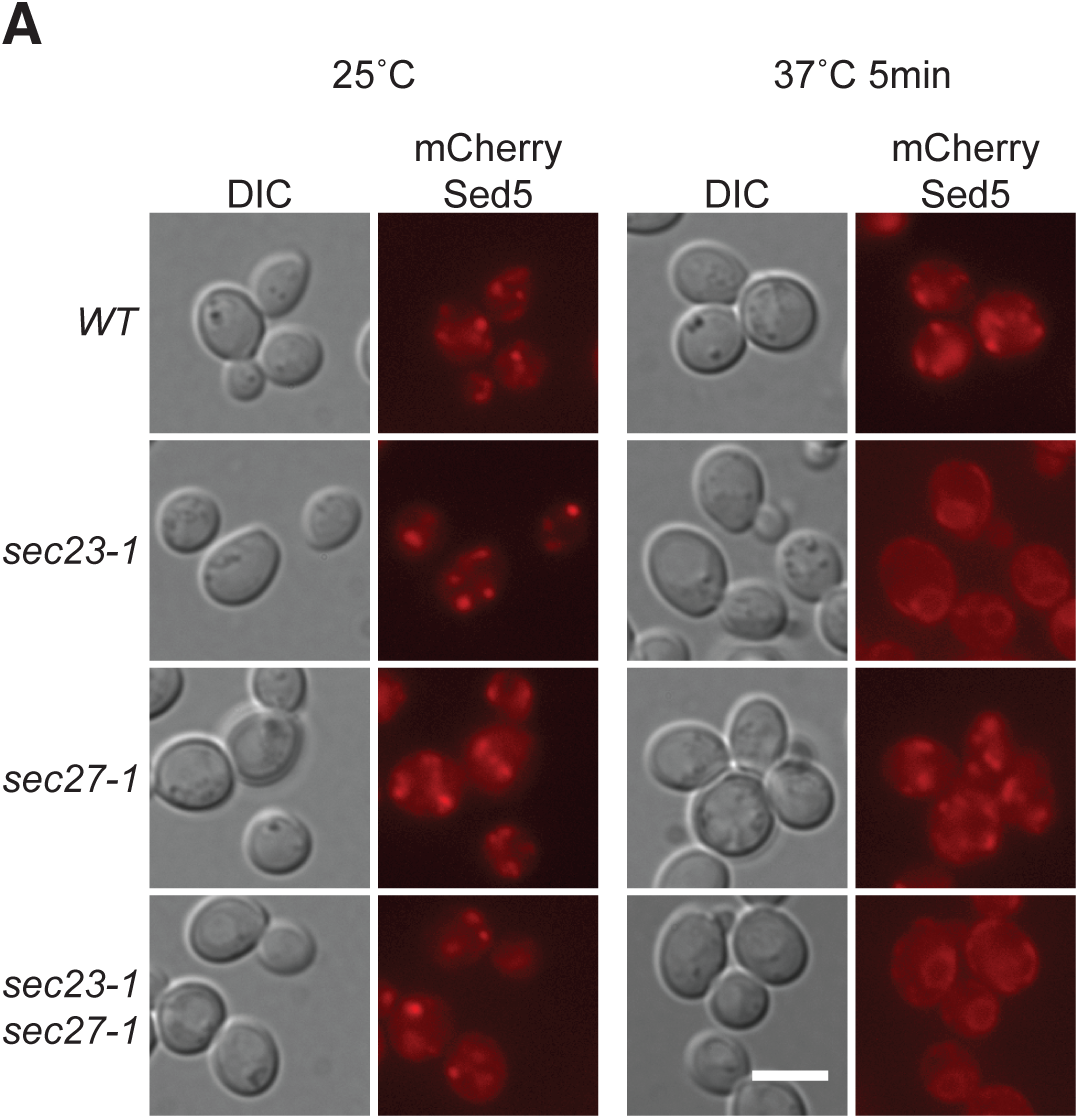
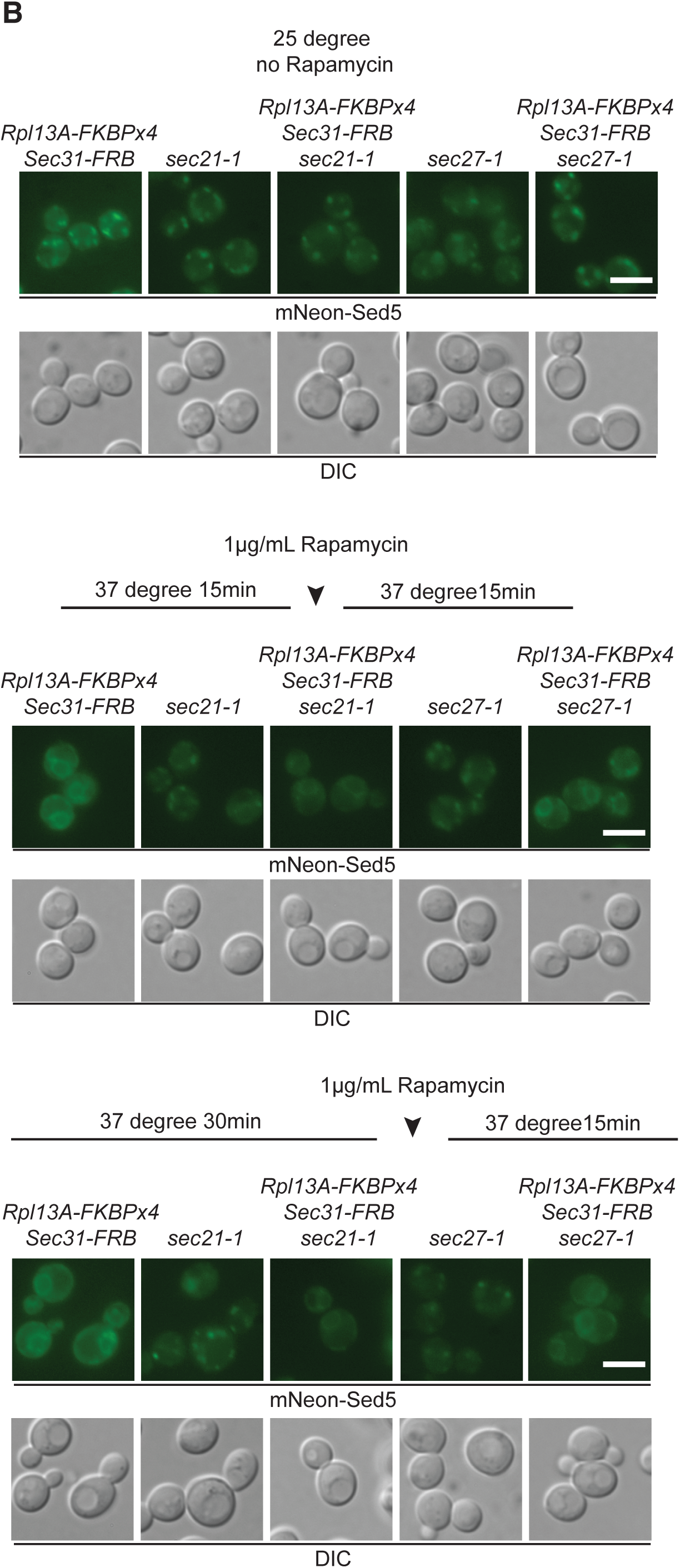
Sed5p’s Golgi – ER trafficking does not require COPI-coatomer function. (A) mCherry Sed5p is still transported from the Golgi to the ER in *sec23-1 sec27-1* cells at the restrictive temperature. (B) mNeon-Sed5p is still transported from the Golgi to the ER in *sec21-1* and *sec27-1* cells at 37°C in which Sec31p was sequestered by rapamycin-based redistribution from the ER to ribosomes (see Material and Methods for details). Scale bar = 5 μm.

### The COPI-binding deficient SED5 mutant (sed5*(KKR)*) shows synthetic genetic interactions with genes involved in intra-Golgi membrane trafficking

To seek additional evidence for a role of the “KKR” motif in the intra-Golgi localization and function of Sed5p we took a genetic approach. *SED5* and *SFT1* have been previously shown to display genetic interactions, and *SFT1* was originally identified as a dosage suppressor of the *sed5-1* temperature-sensitive allele (Banfield *et al*, 1995). Sft1p functions as a t-SNARE together with Sed5p, mediating membrane fusion events within the yeast Golgi (Tsui *et al*, 2001; Parlati *et al*, 2002). If *sed5(KKR)* expressing cells were defective in intra-Golgi trafficking then when combined with mutations in *SFT1* a synthetic growth defect would be expected. While *sed5(KKR)* cells were not temperature-sensitive for growth *sed5(KKR)* s*ft1-6* and *sed5(KKR) sft1-13* cells showed a synthetic temperature-sensitive growth phenotype (Figure 5A). Similarly, while *SED5* could function as a dosage suppressor of the temperature-sensitive growth phenotype of either *sft1-6* or *sft1-13* cells, *sed5(KKR)* could not (Figure 5B).

**Figure 5.**
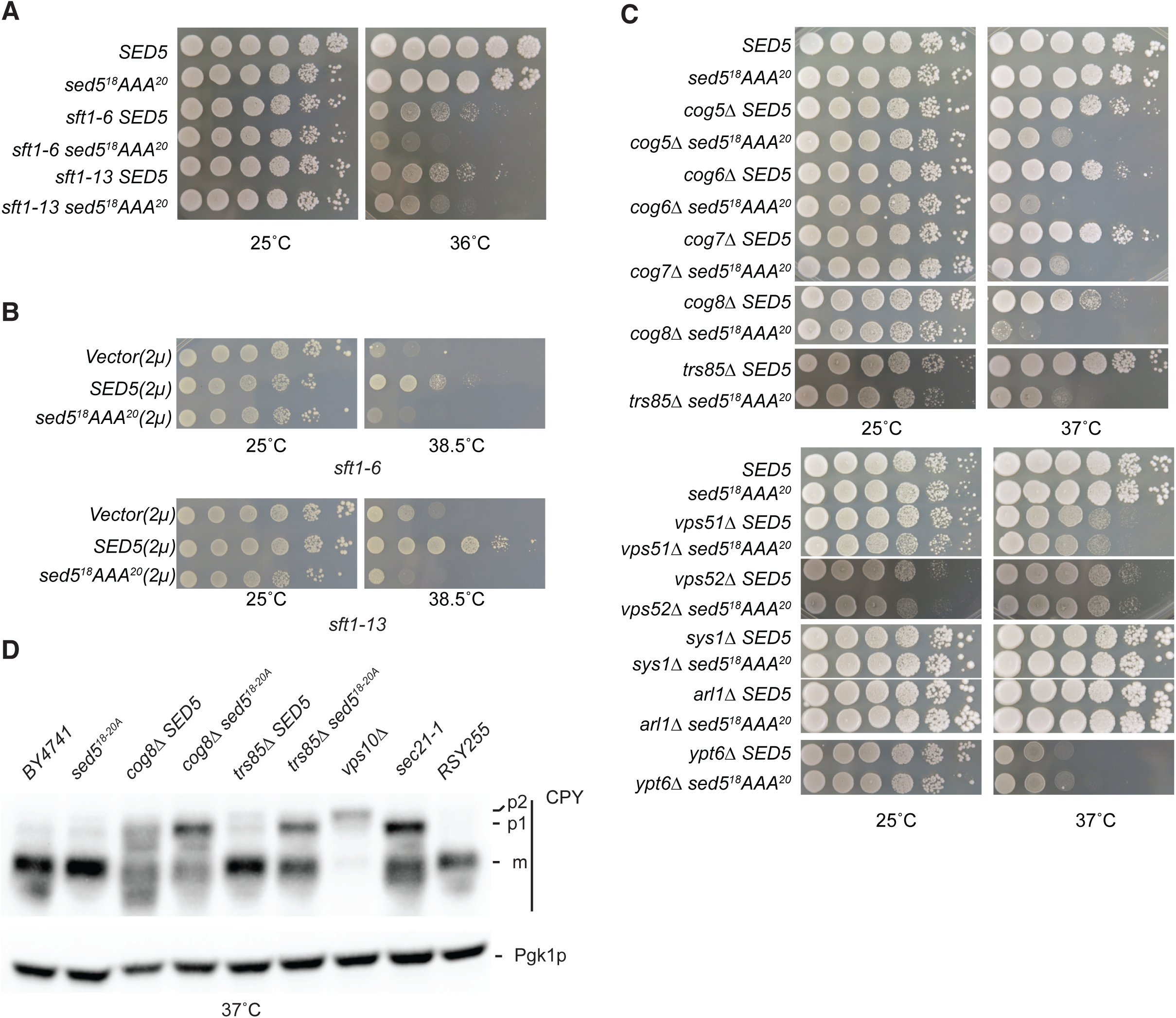
sed5 COPI-coatomer binding deficient mutants show genetic interactions with genes involved in intra-Golgi transport. (A) *sed5* (K18A, K19A, R20A) shows a synthetic temperature-sensitive growth defect with *sft1* mutants. (B) *SED5* but not *sed5* (K18A, K19A, R20A) is a dosage suppressor the temperature-sensitive growth phenotype of *sft1* mutants. (C) *sed5* (K18A, K19A, R20A) shows a synthetic temperature-sensitive growth defect with cells lacking lobe B COG genes, and with cells lacking the TRAPPIII component *TRS85*. No negative genetic interactions are observed between *sed5* (K18A, K19A, R20A) and *arl1*Δ or *sys1*Δ, or *ypt6*Δ or with GARP component encoding genes (*VPS51* and *VPS52*). (A) - (C) 10-fold serial dilutions of the indicated mutant strains were spotted (from left to right) onto nutrient agar plates and incubated at the indicated temperatures for 2 days and then photographed. (D) *sed5*(KKR) cog8*Δ* and sed5(KKR) *trs85Δ* mutants show a synthetic accumulation of carboxypeptidase Y (CPY) trafficking intermediates. BY4741 and RSY255 serve a wildtype controls whereas *vps10Δ* cells accumulate p2 CPY and *sec21-1* cells accumulate p1 CPY. Whole cell extracts of proteins from the indicated strains were resolved by SDS-PAGE and CPY detected by immuno-staining with an anti-CPY antibody. mCPY (mature, vacuolar form), p1 CPY (ER form), p2 CPY (Golgi form). Immuno-staining with an antibody against Pgk1p serves as a gel loading control.

To gain insight into where in the Golgi *sed5(KKR)* localization and /or function was deficient, we generated a series of double mutants in which cells expressing *sed5(KKR)* were combined with deletions of genes whose products are known to be involved in: intra-Golgi transport (COG), Golgi Rab GTPase exchange activity (the TRAPPIII component *TRS85)*, recruitment of GRASP-domain containing Golgins to the Golgi (*SYS1*, *ARL1*), sorting at the TGN (GARP) and the GTPase *YPT6*. When one of either COG5 – 8 was deleted in *sed5(KKR)* cells the double mutants showed a temperature-dependent synthetic growth defect, the same was also true for *sed5(KKR) trs85Δ* cells (Figure 5C). When the transport of carboxypeptidase Y (CPY) was assessed in these temperature-sensitive strains both the *sed5 cog* and *sed5 trs85* double mutants accumulated a processing intermediate of CPY (Figure 5D). In contrast, no synthetic genetic interactions were observed between *sed5(KKR)* and *ARL1* or *SYS1*, with *YPT6* or with GARP component encoding genes (*VPS51* - *VPS52*) (Figure 5C). In sum, these data are consistent with a role for Sed5p’s COPI-binding motif in some facet of intra-Golgi protein trafficking.

### The COPI-coatomer binding motif (^18^KKR^20^) from Sed5p is sufficient to mediate COPI dependent recycling in cells

Both cell biological (Figure 3 and 4) and genetic data (Figure 5) suggested that the “KKR” COPI-coatomer binding motif was required for the intra-Golgi localization and function of Sed5p. To seek more conclusive evidence that the “KKR” motif was necessary and sufficient to direct COPI-mediated trafficking we generated Sed5p-Bet1p chimeras as *in vivo* reporters (Figure 6A). In contrast to Sed5p, the Golgi resident Qc-SNARE Bet1p is found in puncta in wild-type cells but is mislocalized to the vacuole in COPI mutants (Figure 3D). The Golgi-retention signal on Bet1p appears to reside in its N-terminal 41 amino acids as when a form of Bet1p lacking this sequence is expressed in wild-type cells the protein is mislocalized to the vacuole (Figure 6A and B). However, when a Sed5p-Bet1p chimera (Sed5^aa1-41^-Bet1aa^45-117^) was expressed in wild-type yeast cells the protein was primarily found in puncta (although vacuolar fluorescence was also apparent in some cells) (Figure 6B). In contrast, when this construct was expressed in COPI mutants (*sec27-1* or *sec21-1*) it was exclusively localized to the vacuole (Figure 6C) - as was the case in wild-type cells when the ^18^KKR^20^ motif in the Sed5p-Bet1p chimera was replaced by alanine (Figure 6D). In addition, Sed5-Bet-GFP was robustly re-distributed to the ER in *sec23-1* cells following a brief incubation at 37°C (Figure 6E). We therefore concluded that the “KKR“ motif was both necessary and sufficient to function as a *bona fide* COPI sorting signal in yeast cells.

**Figure 6.**
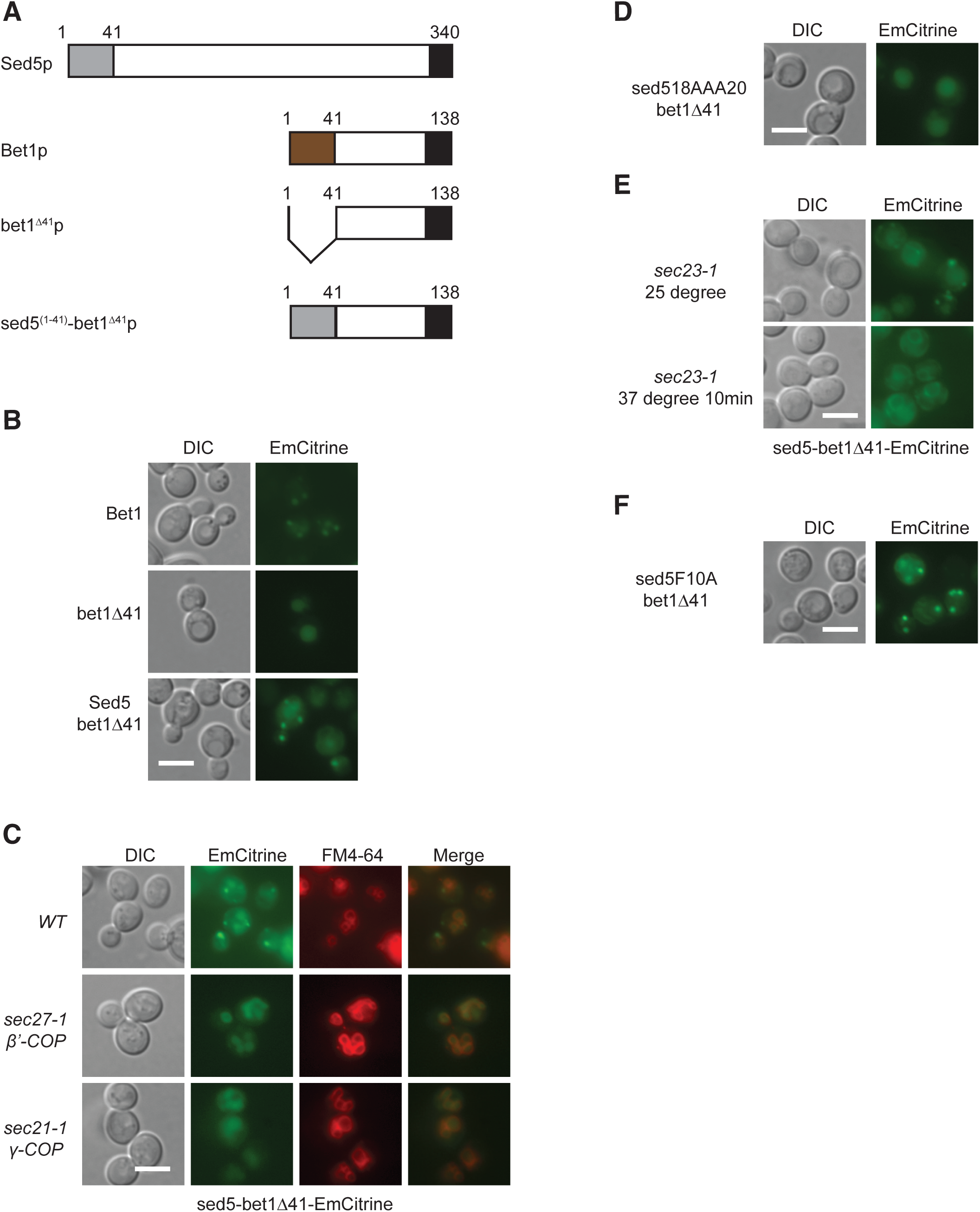
The tribasic COPI-coatomer binding motif from Sed5p is necessary and sufficient to direct COPI-mediated trafficking in yeast cells. (A) Schematic representation of Sed5p / Bet1p constructs used in this study. The N-terminal 41 amino acids of Sed5p contain the COPI-coatomer binding motif (K18, K19, R20). (B) The N-terminal 41 amino acids of Sed5p are sufficient to retain a Sed5-Bet1-emCitrine chimera in the Golgi. (C) The Sed5 ^N41^-Bet1p chimera is mislocalized to the vacuole in COPI mutants. (D) Amino acid substitutions that ablate binding to COPI-coatomer *in vitro* result in mislocalization of the Sed5^N41(AAA)^-Bet1p chimera in wild-type cells. (E) Sed5^N41^-Bet1p chimera can be transported from the Golgi to the ER in *sec23-1* cells. (F) The Sly1p-binding defective form of the Sed5^N41(F10A)^-Bet1p chimera is less prone to vacuolar localization in wild-type cells.

The first 41 amino acids of Sed5p also includes a binding site (aa 5 – 9) for Sed5p’s cognate SM protein – Sly1p, and given that we observed some of the Sed5p-Bet1p chimera in the vacuole, we wondered whether Sly1p might bind to the Sed5p-Bet1p chimera and in doing so occlude coatomer binding to the “KKR” motif. To address this we introduced an additional amino acid substitution into the Sed5p-Bet1p chimera Sed5p F10A (Sed5p(F10A)-Bet1p) that has been shown in biochemical experiments to oblate binding between Sed5p and Sly1p (Peng & Gallwitz, 2004). The Sed5p(F10A)-Bet1p chimera still localized to the Golgi but less vacuolar localization was evident (Figure 6F). In the context of Sed5p, these data are consistent with a model in which Sly1p binding to the N-terminus of Sed5p occludes the Sed5p “KKR” COPI-coatomer binding motif. Given that the *sed5*(KKR) mutant shows genetic interactions with genes that function in intra-Golgi transport, presumably some proportion of Sed5p must be unbound by Sly1p – at least when Sed5p is localized to *cis* distal Golgi cisternae.

### The COPI-coatomer binding motif is occluded when Sly1p is bound to Sed5p

The observation that the “KKR” motif was both necessary and sufficient to confer Golgi localization to Bet1p raised the question of why the motif did not appear to function prevalently on Sed5p. Given the proximity of the ^18^KKR^20^ motif to the high affinity-binding site for Sly1p (aa 5-9), a protein that controls Sed5p’s conformation and thus its accessibility to other SNAREs (Peng & Gallwitz, 2004, 2002; Demircioglu *et al*, 2014; Yamaguchi *et al*, 2002), we next addressed if Sly1p binding might occlude the COPI-coatomer binding site on Sed5p. To address this prospect we conducted biochemical experiments in which GST-Sed5 was immobilized on GSH-beads with increasing amounts of purified bacterially expressed Sly1 followed by the addition of yeast whole cell extracts (Figure 7A). As anticipated, Sly1 was able to compete with coatomer binding to Sed5 (Figure 7A). If the pool of Sed5p in cells is predominantly bound by Sly1p, the “KKR” motif is likely unable to mediate Sed5p binding to COPI-coatomer. Thus, in the context of otherwise wild-type Sed5p, altering the “KKR” motif would be expected to have little to no effect on Sed5p’s Golgi localization - which is precisely what we have observed (Figure 3). In turn, this implies that the interaction between Sly1p and Sed5p is robust, which is borne out in biochemical studies, and by the observation that Sly1p-GFP localizes to the Golgi in cells (Peng & Gallwitz, 2004). Therefore either the interaction between Sly1p and Sed5p is characterized by rapid association and dissociation (that excludes COPI-coatomer binding to “KKR”) or alternatively these proteins remain largely associated one with the other.

**Figure 7.**
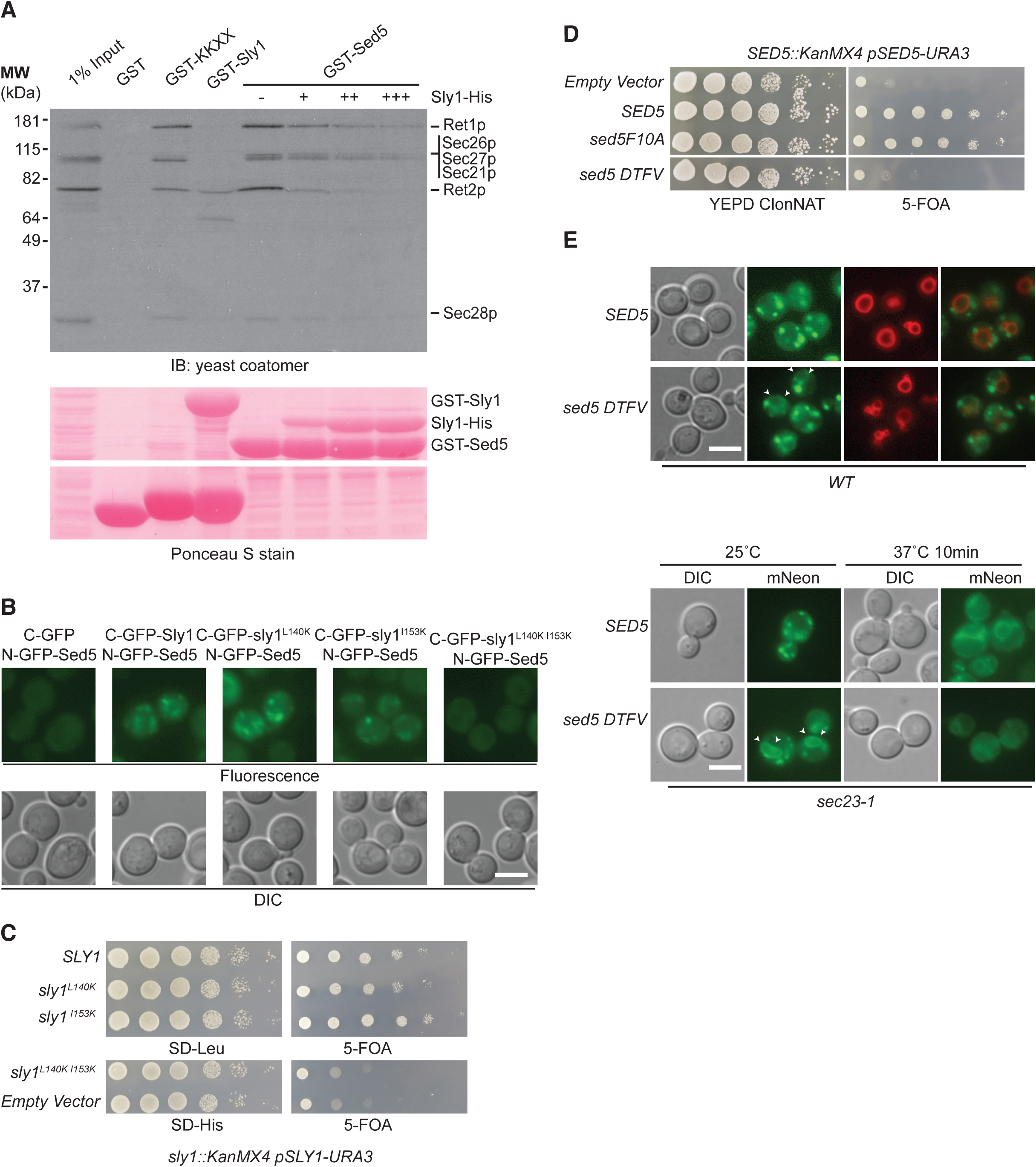
Sly1p binding to Sed5p is essential for growth and affects the association of Sed5p with the COPI and COPII coats. (A) Sly1 competes with COPI-coatomer for binding to Sed5. Increasing amounts of bacterially expressed purified Sly1 were first mixed with Sed5-GST, the complex was then mixed with yeast whole cell lysate. Bound proteins were resolved by SDS-PAGE and COPI-coatomer detected by immuno-staining (top panel). Bottom panel Ponceau S stained membrane used in top panel. (B) Bimolecular fluorescence complementation assay (BiFC) between Sed5p and missense mutants of Sly1p. Scale bar: 5 μm (C) Amino acid substitutions in Sly1p that disrupt association with Sed5p in the BiFC assay result in loss of function of *SLY1*. (D) Combining Amino acid substitutions in Sed5p (D5R, T7A, F10A, V14A) that disrupt binding between Sed5 and Sly1 biochemical assays results in loss of function of *SED5*. (C) and (D) 10-fold serial dilutions of the indicated mutant strains were spotted (from left to right) onto the indicated plates, incubated at 25°C for 2 days and then photographed. (E) mNeon-sed5p (D5R, T7A, F10A, V14A) localizes to puncta and the ER and, can be retrieved to the ER in *sec23-1* cells. Scale bar: 5 μm

Previous reports have indicated that the association between Sed5p and Sly1p is not required for cell growth, as missense mutations in *SED5* (F10A) and *SLY1* (L140K) could support the growth of cells lacking the respective wild-type gene, moreover *sly1*Δ *sed5*Δ cells expressing sly1p (L140K) and sed5p (F10A) were also viable (Peng & Gallwitz, 2004). When we examined the ability of Sly1p to associate with Sed5p using a bimolecular fluorescence complementation (BiFC) assay (Kerppola, 2006) we found that sly1p (L140K) could still bind to Sed5p (Figure 7B), however when we introduced an additional amino acid substitution into Sly1p (which has also been shown to ablate binding to Sed5 *in vitro* (Peng & Gallwitz, 2004) cells expressing this form of sly1p (L140K, I153K) were not viable and sly1p (L140K, I153K) (Figure 7C) did not bind to Sed5p in the BiFC assay (Figure 7B). This finding prompted us to re-examine the results of the biochemical studies with Sed5p and Sly1p (Peng & Gallwitz, 2004). We generated a multiple amino acid substituted form of Sed5p (D5R,T7A,F10A,V14A), hereafter designated sed5p(DTFV), that included residues that singly ablated the interaction between Sed5p and Sly1p *in vitro* (Yamaguchi *et al*, 2002; Peng & Gallwitz, 2004). While *sed5*(DTFV) could not support the growth of cells lacking endogenous *SED5* (Figure 7D), when expressed in wild-type cells, mNeon-sed5p(DTFV) predominantly localized to puncta (Figure 7E) and like wild-type Sed5p this construct was robustly retrieved to the ER in *sec23-1* cells (Figure 7E). We therefore concluded that the association between Sed5p and Sly1p is essential for yeast cell viability. However, Sly1p binding to Sed5p does not contribute to the Golgi localization or the Golgi-ER transport of Sed5p in cells, therefore neither Sly1p-binding to DTFV nor the “KKR” motif appear to substantially contribute to Sed5p’s steady-state Golgi localization

The binding of Sly1p to Ufe1p is also essential for yeast cell growth, as a Sed5p-Ufe1p chimera carrying the DTFV (D5R, T7A, F10A, V14A) substitutions in the Sed5p portion of the fusion protein was unable to support the growth of cells lacking *UFE1* (Figure S2). Curiously, Sly1p does not localize to the ER at steady-state but rather to puncta, and moreover Sly1p cycles between the Golgi and the ER – presumably by virtue of being robustly associated with Sed5p (Li *et al*, 2005). In sum, these observations revealed that binding of Sly1p to the N-terminal motif of both Sed5p and Ufe1p is essential for growth, but the relative affinity, and steady-state binding characteristics of Sly1p to its cognate syntaxins are not equivalent (Yamaguchi *et al*, 2002).

### Sly1p binding is required for robust export of Sed5p from the ER

Unexpectedly sed5p(DTFV) was partially localized to the ER in both wild-type, and in *sec23-1* cells incubated at the permissive temperature (Figure 7E). These findings suggested that Sly1p binding to Sed5p’s DTFV motif might be a prerequisite for the export of Sed5p from the ER. To address this possibility we conducted biochemical experiments in which lysates from yeast cells exclusively expressing a myc-tagged version of Sec24p (Sec24p-9myc) were mixed with increasing amounts of purified Sly1p and Sed5-GST (Figure 8). While Sed5p could bind to Sec24p-9myc in the absence of exogenously added purified Sly1p the relative amount of Sec24p-9myc bound to Sed5-GST increased with increasing amounts of purified Sly1p (Figure 8). By contrast, a mutant form of Sed5p that is defective in binding to both the A and B sites on Sec24p (sed5AB) (Mossessova *et al*, 2003) showed no association with Sec24-9myc across an identical range of Sly1p concentrations (Figure 8). Based on the data presented in Figures 7 and 8 we concluded that Sly1p binding to Sed5p facilitated the export of Sed5p from the ER.

**Figure 8.**
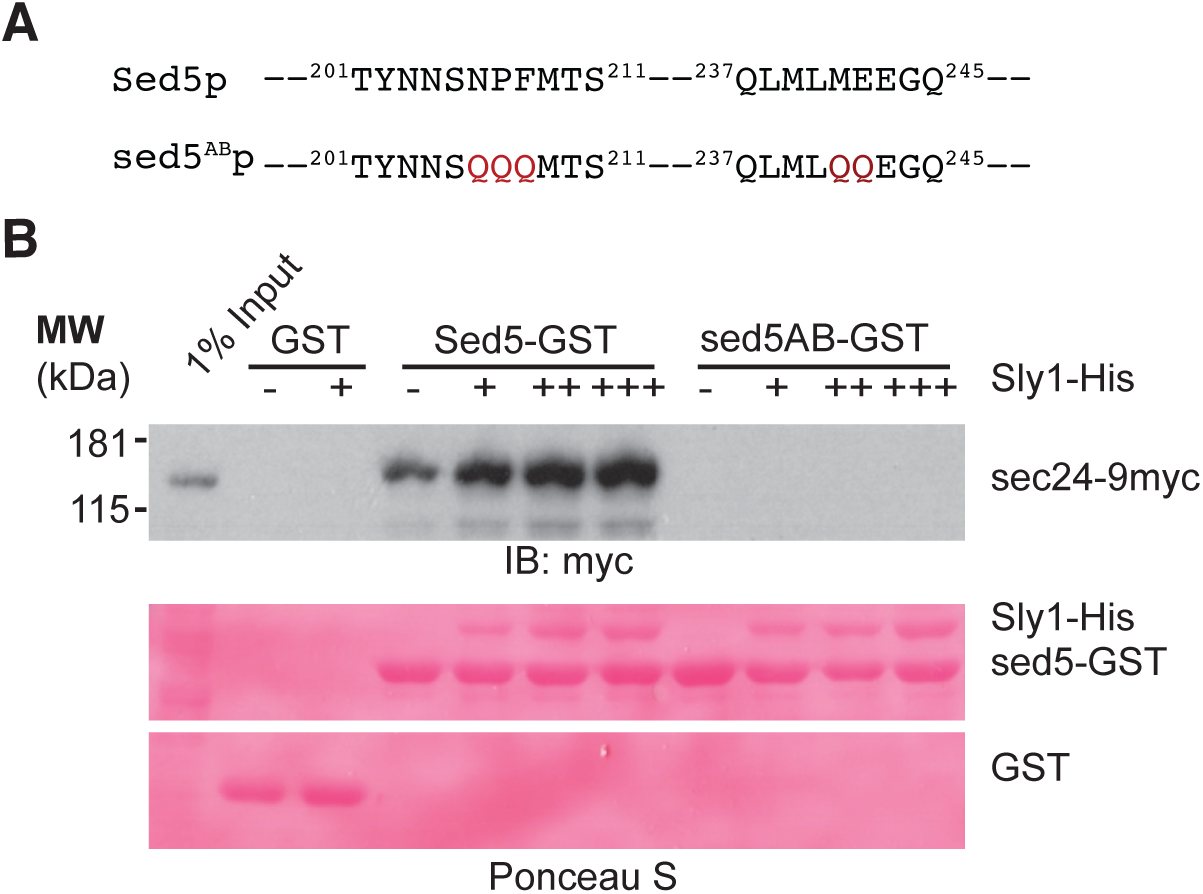
Sly1 binding to Sed5 enhances the association between COPII and Sed5 in vitro. (A) Depiction of the amino acid substitutions in the COPII-binding deficient mutant of sed5p (sed5p^AB^) used in these experiments. (B) Increasing amounts of purified bacterially expressed Sly1 were mixed with GST, Sed5-GST or sed5^AB^-GST after which the complex was incubated with whole cell extracts from cells expressing Sec24-9myc. Bound proteins were visualized following SDS-PAGE with Ponsceau S (bottom panel) and immuno-stained with an anti-myc antibody.

### Missense mutations in the SNARE-motif disrupt the Golgi localization of Sed5p

In contrast to other Golgi SNAREs, mNeon-Sed5p remained in puncta in COPI-coatomer mutants (Figure 3). While these data are consistent with the rather modest contribution of the KKR motif to Sed5p’s Golgi localization, they also suggested that COPI function *per se* is not required for Sed5p’s steady-state localization to the Golgi. Similarly, the amino acid sequence of the transmembrane domain was not sufficient to account for Sed5p’s Golgi localization (Figure 1). In addition, Golgi localization of Sed5p was not mediated by a combination of the KKR motif and the amino acid sequence of the TMD as sed5 (KKR-AAA)-Sso1 (TMD) was not mislocalized (Figure S3), nor by the binding of Sly1p to Sed5p (Figure 7). Moreover, a form of sed5p deficient in both Sly1p and COPI-coatomer binding (mNeon-sed5p(DTFV+KKR)) was also not mislocalized (Figure S4).

What then accounts for Sed5p’s Golgi localization? To delve deeper into this question we took a genetic approach based on the functional relationships between Sed5p, Sly1p and Ufe1p (Li *et al*, 2005). The SM protein Sly1p binds to an N-terminal motif found in both Sed5p as well as Ufe1p (Yamaguchi *et al*, 2002), and we show here that the binding of Sly1p to both proteins is essential for Sly1p (Figure 7) as well as for Sed5p and Ufe1p function (Figure 7). However, at steady-state Sly1p is predominantly found on Golgi membranes (Li *et al*, 2005), and we therefore surmised that the binding between Sed5p and Sly1p was more robust than that between Sly1p and Ufe1p in cells – a higher binding affinity of Sly1p for Sed5p has also been described in yeast two-hybrid experiments (Yamaguchi *et al*, 2002). Indeed, the steady-state localization of Sly1p is partially redistributed to the ER in cells expressing an N-terminal Sed5p-Ufe1p chimera (Figure S5).

These observations suggested that Sly1p might bind to Ufe1p transiently, or alternatively, in *trans* or in *cis* when the complex of Sed5p and Sly1p cycled through the ER. To test this directly we asked if a Sly1p-Sed5p fusion protein was functional. Although, the Sly1p-EmCitrine-Sed5p construct could not support the growth of cells lacking *SLY1* (Figure S6A) *or SED5 and SLY1* (Figure 9C), when a missense mutation that weakens the interaction between Sly1p and Sed5p *in vitro* was introduced into the Sed5p portion of the chimera the resulting protein ((Sly1p-EmCitrine-sed5p(F10A)) could support the growth of cells lacking both *SLY1* and *SED5* (Figure 9C) and the fusion protein was localized to puncta (Figure 9B). Interestingly, even when Sly1p and Sed5p were covalently linked, the N-terminal 21 amino acids of Sed5p were required in order for the chimera to support the growth of cells lacking *SED5* but this construct could support the growth of *sly1Δ* cells (Figure S6B).

**Figure 9.**
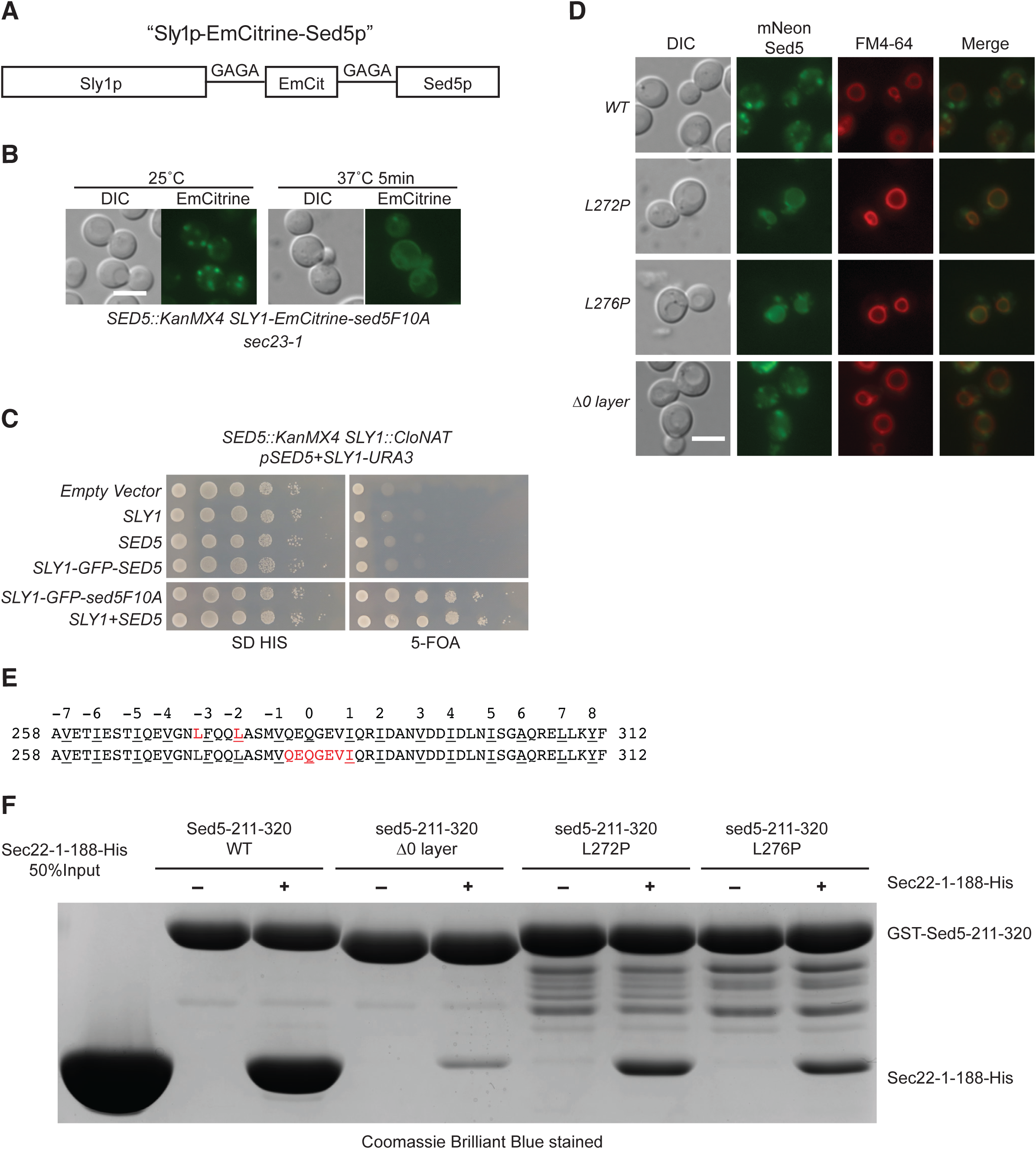
Missense mutations in the SNARE-motif disrupt the Golgi localization of Sed5p. (A) Schematic representation of the Sly1p-emCitrine-Sed5p fusion protein. (B) Sly1p-EmCitrine-sed5p^F10A^ localizes to the Golgi and can be transported from the Golgi – ER in *sec23-1* cells. Scale bar: 5 μm (C) Sly1p-EmCitrine-sed5p^F10A^ can support the essential functions of both *SLY1* and *SED5*. 10-fold serial dilutions of the indicated mutant strains were spotted (from left to right) onto the indicated plates, incubated at 25°C for 2 days and then photographed. (D) Missense mutations in the SNARE-motif result in mislocalization of mNeon-Sed5p to the limiting membrane of the vacuole. Scale bar: 5 μm (E) Amino acid sequence alignment of the SNARE-motif of Sed5p. The missense mutations that result in mislocalization of Sed5p to the limiting membrane of the vacuole (above) and the amino acids that define the zero-layer (below) are in red font. The numbering above the alignments denotes the heptad repeats and layers of the SNARE-motif. (F) The missense mutations that result in mislocalization of mNeon-Sed5p to the limiting membrane of the vacuole do not disrupt interactions between the Sed5 and Sec22 *in vitro*.

Like Sed5p, Sly1p-EmCitrine-sed5p(F10A) was robustly redistributed to the ER in *sec23-1* cells following a short incubation at the non-permissive temperature (Figure 9B). We therefore reasoned that in cells expressing this Sly1p-Sed5p chimera as their sole source of both Sly1p and Sed5p, Ufe1p could only associate with Sly1p from the chimeric protein (i.e. Sly1p-EmCitrine-sed5p(F10A). Consequently then any mutation in within the *SED5* portion of the chimera that disrupted Sed5p’s Golgi retention / Golgi - ER transport should be lethal to cells lacking endogenous Sly1p. Similarly, if the sole source of the cell’s Sly1p were covalently linked to Sed5p, mislocalization of such a chimera would prevent Sly1p from binding to endogenous Sed5p and Ufe1p. On the basis of these premises we designed a genetic screen using the Sly1-EmCitrine-Sed5p construct as a means by which to identify mutations in *SED5* that disrupted Sed5p’s Golgi - ER retrieval / Golgi retention.

To identify mutants that were defective in the retention of Sed5p in the Golgi we generated a collection of random mutations in *SED5* using error-prone PCR (Muhlrad *et al*, 1992) and introduced them into the *SED5* portion of the *SLY1*-EmCitrine-*SED5* construct. The mutant collection was then screened to identify alleles that could not support the growth of cells in which the endogenous *SLY1* promoter had been replaced with the regulatable *GAL4* promoter (Table 1). Transformants were initially selected on media containing galactose, and then replicated onto media containing glucose to identify retention-defective mutants. Using this strategy we identified two alleles of *SED5* that could not support the growth of cells lacking the *SLY1* gene. One allele contained a missense mutation that resulted in a substitution of leucine with proline at position 272 (L272P) of Sed5p while the other allele contained a missense mutation that resulted in a substitution of leucine with proline at position 276 (L276P) of Sed5p. Fluorescence microscopy revealed that both mutant proteins (mNeon-sed5p(L272P) and mNeon-sed5p(L276P) were mislocalized to the limiting membrane of the vacuole, as judged by colocalization with FM4-64 dye. (Figure 9D).

The L272 and L276 residues are located in the SNARE-motif of Sed5p (Figure 9E) and substitution of leucine with proline might therefore be expected to perturb the formation of any intermolecular helices required for SNARE complex formation. Thus in principle, the loss of Golgi retention observed for the *sed5*L272P and *sed5*L276P mutants could be related to an inability of the mutant proteins to form SNARE complexes. To examine the capacity of the sed5p mutants to bind SNAREs we conducted binding assays between Sed5p and the various sed5p mutants with the canonical SNARE binding-partner Sec22p (Figure 9F). When a GST-tagged form of the SNARE-motif from wild-type Sed5p was mixed with purified recombinant Sec22p robust binding was observed (Figure 9F). This binding was reduced when GST-sed5p(L272P) or GST-sed5p(L276P) was used in place of the wild-type protein (Figure 9F). However, deletion of the zero-layer residue of Sed5p (sed5Δ0, Figure 9 E and F), which should impede SNARE-complex formation (Fasshauer *et al*, 1998; Graf *et al*, 2005), bound less well to Sec22p than either of the Leu to Pro substitution mutants (Figure 9F). Moreover, unlike sed5p(L272P) and sed5p(L276P), which localized to the limiting membrane of the vacuole sed5pΔ0 remained in puncta (Figure 9D). Thus failure to form SNARE complexes *per se* did not account for the loss of Golgi localization observed for either sed5p(L272P) or sed5p(L276P).

### Sed5p’s Golgi retention is conformation-dependent

The SNARE-motif of Sed5p is known to mediate SNARE-SNARE interactions as well as mediate an intra-molecular interaction between the SNARE-motif and the Habc domain of Sed5p (Demircioglu *et al*, 2014). Our findings (Figure 9) suggested defects in SNARE complex formation did not account for the mislocalization of either sed5p(L272P) or sed5p(L276P) to the limiting membrane of the vacuole. We therefore next asked, using an *in vitro* binding assay, whether sed5p(L272P) or sed5p(L276P) mutant proteins were defective in binding to the Habc domain of Sed5p (Figure 10).

**Figure 10.**
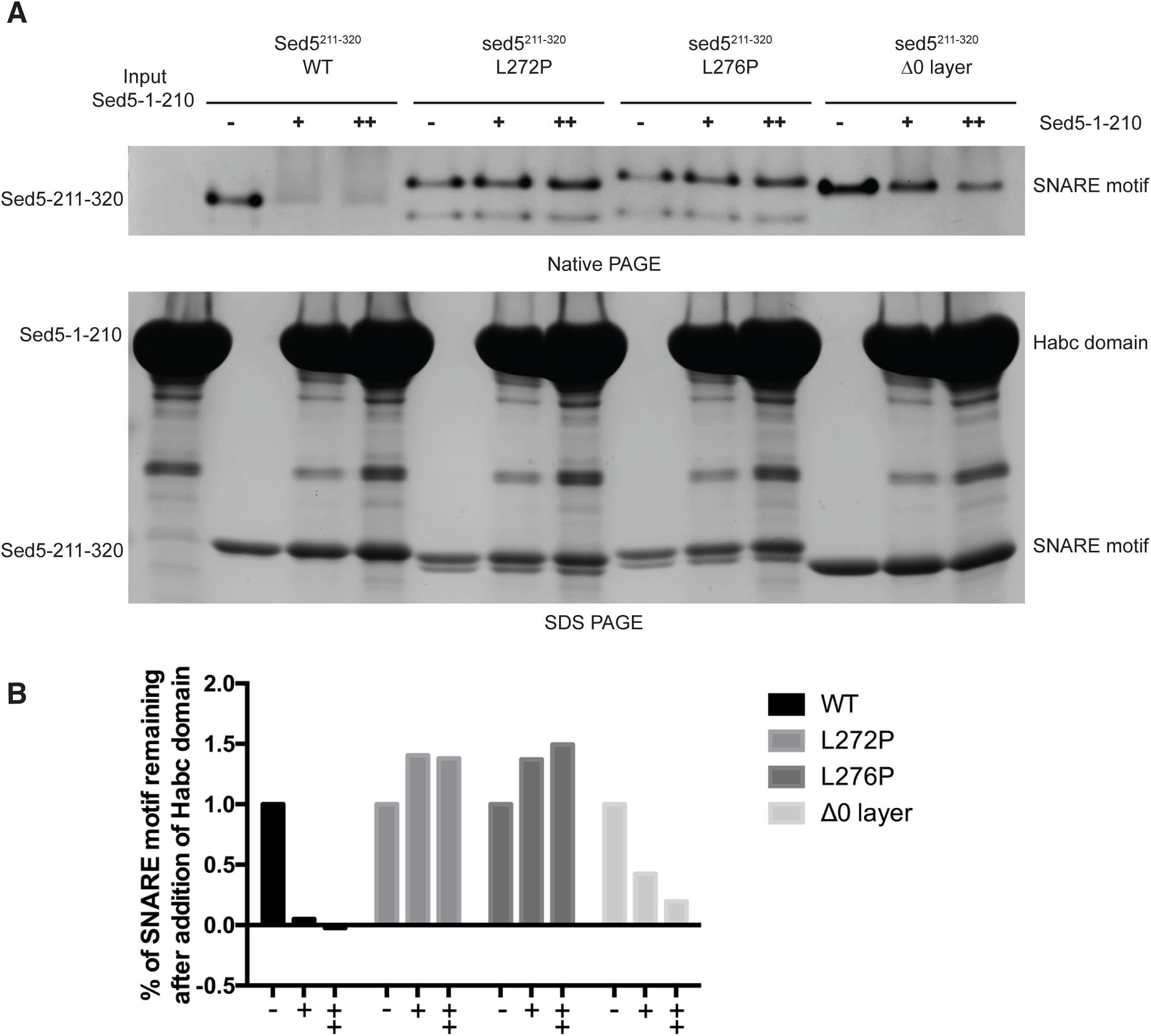
The missense mutations that result in mislocalization of Sed5p to the limiting membrane of the vacuole disrupt interactions between the Habc domain and SNARE-motif in vitro. (A) Increasing amounts of bacterially expressed purified Habc domain (amino acids 1-210) were mixed with bacterially expressed purified SNARE-motif (amino acids 211-320). Following incubation binary complex formation was detected by native PAGE and staining with Coomassie Brilliant Blue (upper panel). Recombinant proteins used in these experiments were resolved by SDS-PAGE and stained with Coomassie Brilliant Blue (bottom panel). Association between the Habc domain and the SNARE-motif results in a band shift of the SNARE-motif upon native PAGE (upper panel). (B) Quantification of the binding data shown in (A). For each set of mixing experiments (WT, L272P, L276P and Δ0) the “-“ sample was arbitrarily set to 100% and the remaining two data points (+ and ++) represent a ratio relative the “-“ sample. Data was obtained using ImageJ software.

Sed5p’s Habc domain (1-210) and SNARE-motif (211-320) have been shown to interact with each other *in vitro* and this interaction could be resolved by native (non-denaturing) polyacrylamide gel electrophoresis (PAGE) (Demircioglu *et al*, 2014). We therefore used this assay to assess interactions between sed5p(L272P) and sed5p(L276P) with the Sed5p Habc domain (Figure 10). We incubated bacterially expressed wild-type Habc domain with wild-type, L272P, L276P and Δ0-layer versions of the Sed5p SNARE-motif and measured their ability to interact by non-denaturing PAGE (Figure 10). As previously reported (Demircioglu *et al*, 2014) the Habc domain peptide cannot be visualized on native-PAGE gels because of its positive charge, however the SNARE-motif is visible, and a reduction in the amount of the SNARE-motif resolved on native PAGE gels correlates well with the formation of a binary complex between the Habc and SNARE-motif domains (Demircioglu *et al*, 2014). While the wild-type Sed5p SNARE-motif and Habc domain readily formed a complex, as judged by the apparent absence of the SNARE-motif on the non-denaturing PAGE gel, neither the L272P nor L276P SNARE-motifs bound to the wild-type Habc domain (Figure 10A and B). By contrast, the Δ0-layer sed5p mutant could still form the Habc and SNARE-motif complex, although to a much lesser extent than wild-type Sed5p (Figure 10A and B). We therefore concluded that the mislocalization of sed5p(L272P) and sed5p(L276P) to the limiting membrane of the vacuole most likely resulted from the inability of these proteins to adopt a folded-back conformation in which the SNARE-motif bound the Habc domain, rather than from a defect in SNARE-complex formation. Thus, Sed5p’s Golgi retention would appear to be primarily conformation-based, rather than Sly1p-dependent, COPI-dependent or TMD-based.

## DISCUSSION

In this study we identify multiple mechanisms that contribute to the localization of Sed5p to the Golgi in budding yeast cells. We have re-investigated the role of the transmembrane domain (TMD) and confirmed that the TMD is not the only feature of Sed5p that contributes to its Golgi localization (Figure 1). We have also identified and characterized a tri-basic COPI-coatomer binding motif located in the N-terminus of Sed5p (^18^KKR^20^) (Figure 2). The Sed5p and Syn5L (long form of mammalian syntaxin 5) tribasic motifs are similar to the ones found in the N-terminus of Vps74p and its mammalian homologs - GOLPH3 and GOLPH3L (Figure 2; Tu et al., 2012). Like the tribasic motif in Sed5p and Syn5L, the Vps74p motif also mediates direct binding to COPI-coatomer (Tu et al., 2012). Surprisingly amino acid substitutions to the (^18^KKR^20^) motif in Sed5p did not result in loss of function, mislocalization of the protein nor impact its robust Golgi – ER retrieval (Figure 3, 4, S3 and S4). Consistent with these observations we found that unlike Bet1p (a Sed5p-binding Golgi Qc-SNARE) and the Golgi resident mannosyltransferases: Mnn9p, Och1p and Ktr3p – Sed5p remained in Golgi-like puncta in temperature-sensitive COPI mutants (Figure 4 and Figure S1). These data were perplexing as they did not provide an explanation for the role of the COPI-binding motif on Sed5p nor explain mechanistically how Sed5p was transported from the Golgi to the ER in cells (Figures 3 and 4). However, these findings are consistent with observations made by Kurokawa et al, (2014) using SCLIM in which they show direct contact between the *cis* Golgi (the Sed5p compartment) and ER exit sites (ERES) during which cargo (the plasma membrane transmembrane protein Alx2-GFP) was transferred directly from ERES to the *cis* Golgi (Kurokawa et al, 2014). It appears that the retention of Sed5p in the Golgi as well as the transport of Sed5p between the *cis* Golgi and the ER proceeds independently of COPI-coatomer function.

The Sed5p (^18^KKR^20^) motif does however satisfy the characteristics of a *bona fide* COPI-coatomer sorting signal when appended to another protein (Figure 6), and genetic interaction studies revealed a role for the (^18^KKR^20^) motif in Sed5p in intra-Golgi function (Figure 5). Additionally, in cells expressing a loss-of-function mutant form of Sed5p that does not bind to Sly1p (sed5p(DTFV)) (Figure 7) the protein localized to the ER and Golgi, but when the (^18^KKR^20^) motif was substituted with alanine this protein was localized predominantly to puncta – observations that are consistent with the (^18^KKR^20^) motif being accessible to COPI-coatomer but only if Sly1p is not bound to the N-terminus of Sed5p (Figures 7 and S4). Thus, neither Sly1p-binding to the N-terminus of Sed5p nor the tribasic (^18^KKR^20^) motif could account for Sed5p’s steady-state localization to the Golgi or the trafficking of Sed5p between the Golgi and ER.

Our observations that the ^18^KKR^20^ motif could function as a COPI-coatomer sorting signal if Sly1p was not bound to Sed5p, and that cells expressing Sed5p in which the (^18^KKR^20^) motif has been substituted with alanine show genetic interactions with genes encoding intra-Golgi transport machinery (Figure 5) - as well as alteration in the protein’s Golgi distribution (Figure 3) - implied that there was indeed a physiological role for this motif. We can envision a number of scenarios for the function of the Sed5p (^18^KKR^20^) motif. For example, the Sed5p (^18^KKR^20^) motif may function to prevent Sed5p that is not bound to Sly1p from leaving the Golgi for more distal endomembrane compartments where Sed5p might otherwise engage non-canonical SNAREs. The KKR motif could function to ensure that Sed5p that is not bound to Sly1p remains primarily in the *cis*-Golgi. In addition / or alternatively, the Sed5p (^18^KKR^20^) motif might function as a failsafe, ensuring that Sed5p that has lost Sly1p is returned to a locale (the cis Golgi or ER) where Sly1p can be re-acquired. Lastly, Sed5p may have an important but non-essential role in the Golgi that does not require association with Sly1p, and given our genetic interaction data (Figure 5) this would most likely occur in the medial / late Golgi.

The functional significance of the association of Sly1p with Sed5p with regard to membrane fusion has been well studied (Lobingier *et al*, 2014). Here we identified an additional role for Sly1p in promoting the robust ER export of Sed5p. In cells the ER export of sed5p(DTFV) is delayed in the COPII mutant *sec23-1* at the permissive temperature (Figure 7E) and *in vitro* Sly1p increases the binding affinity of Sed5p for Sec24p a COPII component to which Sed5p binds directly (Mossessova *et al.*, 2003; Mancias and Goldberg, 2007) (Figure 8). These observations may help to reconcile contrasting data on the mechanism by which Sed5p is preferentially packaged into COPII vesicles. According to structural and biochemical studies, the open form of Sed5p should be preferably packaged into COPII vesicles, as the Sec24p binding-motif (Sed5p(“NPY”)) is occluded when the protein adopts a folded-back conformation (Mossessova *et al*, 2003). In yeast cells however it appears that Sed5p is not sorted into COPII vesicles as part of a cognate SNARE complex. Sed5p bearing missense mutations that disrupt binding to the Sec24 A-site were excluded from COPII vesicles, whereas Sed5p’s cognate SNAREs - Bet1p and Bos1p - were not (Miller *et al*, 2005). If the primary mechanism by which Sed5p is selectively packaged into COPII vesicles requires an accessible “NPY” motif for binding to the Sec24p A-site then Sed5p should be in the open conformation – but not in a complex with its cognate SNAREs. Formation of the folded-back conformation of Sed5p *in vitro* can occur in the absence of supplementary proteins, and others have shown that the addition of Sly1p could disrupt this folded-back conformation. The authors’ speculated that this in turn accelerated the formation of Sed5p-containing SNARE complexes - presumably by mediating the transition of Sed5p from the closed to open conformation (Demircioglu *et al*. 2014). Perhaps the propensity of Sly1p to alter the balance of the closed to open form of Sed5p together with mass action effects brought about by the high local concentration of Sec24p at ER export sites can satisfy the requirement that Sed5p be sorted in an open conformation.

To identify the mechanism of Sed5p’s localization to the Golgi we exploited the observations that the association between Sed5p and Sly1p, as well as between Sly1p and Ufe1p, were essential for growth (Figure 7, Figure S6). Based on these findings we surmised that if the Sly1-mNeon-sed5p(F10A) fusion protein were mislocalized in *sly1Δ* cells it would be lethal, as Sly1p (Sly1p-mNeon-sed5p(F10A)) would not be available to bind to either endogenous Sed5p or to Ufe1p. We identified several missense mutations within the *SED5* portion of Sly1p-mNeon-sed5p(F10A) that were lethal in cells conditionally lacking endogenous *SLY1* and that also resulted in mislocalization of the mNeon-sed5p to the limiting membrane of the vacuole in cells (Figure 9). Biochemical experiments established that two of the missense mutants we characterized (L272P and L276P) did not ablate SNARE complex formation, but rather were defective in the formation of the folded-back conformation of Sed5p - in which the N-terminal Habc domain contacts the SNARE-motif (Figure 10). It is therefore plausible that the folded-back conformation is also an inherent property of Sed5p in cells. Indeed the Golgi – ER trafficking of Sed5p is independent of Sly1p binding to the N-terminus of Sed5p (Figure 7E), and if our *in vitro* findings recapitulate the physiological scenario then Sly1p binding to Sed5p seems unlikely to play a direct role in the conformation-dependent Golgi retention and Golgi - ER transport of Sed5p.

Two additional interesting findings were made during these studies. Firstly, the Sly1p-mNeon-Sed5p chimera could support the growth *sed5Δ* cells but not *sly1Δ* cells (Figure 4S). We surmised that this was the result of an increase in the association of Sly1p with Sed5p arising from an intra-molecular interaction rather than an inter-molecular interaction (that would occur in wild-type cells) and that this in turn impeded the association of Sly1p (Sly1p-mNeon-Sed5p) with Ufe1p. Secondly, a Sly1p-mNeon-Sed5p in which the N-terminal 21 amino acids had been deleted (Sly1p-mNeon-sed5pΔN21, Figure S6) whilst supporting the growth of *sly1Δ* cells could not support the growth of *sed5Δ* cells. Apparently, the first 21 amino acids of Sed5p do not simply function to facilitate the binding of Sly1p to Sed5p but must also be important for the mode of binding – perhaps by inducing a conformational change in Sly1p (Baker et al., 2015).

One striking observation made in this study concerns an unanticipated aspect of the functional relationship between Sly1p and it’s ER-localized cognate syntaxin Ufe1p (Lewis et al., 1997). A previous study reported that when Ufe1p was not bound to Sly1p, Ufe1p was ubiquitinated and subsequently robustly degraded (Braun and Jentsch, 2007). Moreover, Ufe1p (but not Sed5p) degradation is particularly sensitive to the level of Sly1p in cells (Braun and Jentsch, 2007). One interpretation of this observation is that Sly1p and Ufe1p need to be constitutively bound to one another, and that ubiquitination of unbound Ufe1p functions as a failsafe by scavenging this pool of the protein in cells. However, our findings that Golgi-localized Sly1p-Sed5p fusion proteins (Sly1p-mNeon-sed5p(F10A) and Sly1p-mNeon-sed5pΔN21) can support the growth of cells lacking endogenous Sly1p suggest that the association between Sly1p and Ufe1p may ordinarily be transient. Furthermore, the propensity for Ufe1p to be degraded in the absence of Sly1p binding suggests that contacts between Ufe1p and Sly1p (that is bound to Sed5p) must frequent. Given that Sly1p is localized to the Golgi at steady-state (Li et al 2005), and that Sed5p’s Golgi localization and Golgi-ER transport is COPI-independent is consistent with the observation that the *cis* Golgi directly contacts the ER (Kurokawa et al, 2014)

Whilst the mechanism of retention mediated by the transmembrane domain of SNAREs and other type II membrane proteins have been proposed (Sharpe et al, 2010), and the role of sorting motifs in mediating coat interactions are well established (Aridor and Traub, 2002; Arakel and Schwappach, 2018) less is known about how the folded-back conformation of SNAREs, and syntaxins in particular, would facilitate organelle retention / membrane trafficking (Tochio et al., 2001; Pryor et al., 2008). We have shown here that Sed5p’s Golgi – ER transport is independent of COPI function whereas it’s cognate binding partners, Bet1p, Bos1p and Sec22p are dependent on COPI-coatomer for their Golgi localization / Golgi – ER transport (Ballensiefen et al., 1998: Ossipov et al., 1999). Taken together, these data suggest that Sed5p does not cycle between the Golgi and the ER in a SNARE-complex. How then would the folded-back conformation of Sed5p mediate interactions with the ER and in particularly in the vicinity of ER exit sites (ERES) or ER acceptor sites (ERAS)? The apparent proximity between ERES and ERAS in yeast cells may be relevant in this regard (Zink et al., 2009). In the folded-back conformation Sed5p may bind to tethering factors on ER membranes – such as components of the DSL complex (Schroter et al., 2016; Koike and Jahn, 2019), or perhaps to one or more COPII proteins. Of course conformational binding epitopes on the closed form of Sed5p could also mediate associations with other proteins on ER membranes – such as the ER localized SNARE proteins.

In summary we have identified and characterized three features of Sed5p that contribute to the protein’s steady-state localization to the Golgi: the amino acid composition of the transmembrane domain, the COPI-coatomer binding motif (^18^KKR^20^), and the folded-back conformation of the protein. While the amino acid composition of the transmembrane domain and COPI-coatomer binding motif are invariant features of the protein – the conformation of Sed5p is presumably subject to change. The predominate mechanism of Golgi retention seemingly involves COPI-coatomer independent iterative contacts between the *cis* Golgi and the ER mediated by the folded-back conformation of Sed5p. This process would be expected to maintain the steady-state distribution of Sed5p in the *cis* Golgi whilst leaving other membrane constituents of the cis Golgi free to move to more distal compartments or to recycle between the Golgi and ER in a COPI-dependent manner. In any case, the multifaceted localization mechanisms adopted by Sed5p underscore the central role that this syntaxin family member plays in the Golgi and our work now offers a more complete picture of how this important SNARE protein is localized.

## MATERIALS AND METHODS

### Materials

Glutathione Sepharose 4B was purchased from GE healthcare. Ni-NTA agarose was purchased from QIAGEN. Strep-Tactin beads were purchased from IBA GmbH. Protease inhibitors [ethylenediamine tetra acetic acid (EDTA)-free complete protease inhibitor cocktail, Pefabloc SC (AEBSF)] were purchased from Roche. FM™ 4-64 Dye (N-(3-Triethylammoniumpropyl)-4-(6-(4-(Diethylamino) Phenyl) Hexatrienyl) Pyridinium Dibromide) was purchased from Invitrogen. Rapamycin was purchased from LC Laboratories. The anti-rat β-COP (496 – 513) antibody was purchased from Calbiochem. The anti-rabbit IgG horseradish peroxidase-conjugated antibody, anti-mouse IgG horseradish peroxidase-conjugated antibody, concanavalin A (C7275) and Ponceau S were purchased from Sigma. The DC Protein Assay Kit was purchased from Bio-Rad. The anti-GFP antibody was purchased from Clontech. The anti-CPY and anti-Pgk1 antibodies were from Molecular Probes. The anti-c-Myc antibody was purchased from Roche. Restriction enzymes and Gibson assemble master mix was purchased from New England Biolabs. The anti-yeast coatomer antibody was kindly provided by Prof. Anne Spang (University of Basel).

### Methods

#### Expression and purification of recombinant proteins

Recombinant proteins were expressed in Escherichia coli BL21 (DE3) cells. Protein expression was induced when cell cultures reached an OD_600_ of ∼0.8 by the addition of 1 mM IPTG and incubation at 18°C overnight. Following protein induction, bacterial cultures were harvested and resuspended in 10 mL Buffer A (25 mM Tris-HCl, pH 7.5, 400 mM KCl, 10% glycerol, 4% Triton X-100) supplemented with 1× EDTA-free complete protease inhibitor, 2mM DTT and 1 mM Pefabloc. Cells were lysed using a constant cell disruption system (Constant System) at 25 kpsi with a pre-cooled chamber. Bacterial lysates were cleared by centrifugation at 16000×g for 10min at 4°C.

For purification of Sed5 SNARE motif, cell lysate expressing GST-fusion protein were first incubated with Glutathione sepharose beads. After extensive washing, thrombin cleavage were conducted in standard buffer (20 mM Tris-HCl, pH 7.4, 200mM NaCl) with 0.03 U/μl of thrombin (Sigma-Aldrich) at 4°C for 6 hours.

For purification of His-tagged recombinant proteins, cell lysates were first incubated with Ni-NTA Agarose beads in the standard buffer in presence of 10mM imidazole. After washing with standard buffer containing 50mM imidazole, elution was conducted using standard buffer containing 300mM imidazole. Desalting was conducted using PD-10 column (GE healthcare) following manufacture’s protocol.

#### Wide field fluorescence microscopy

Yeast cells were grown in appropriate selective medium at 25°C to mid-log phase (OD_660_ = 0.6 - 0.8). Cells were harvest by centrifugation and washed by ddH_2_O for once and then resuspended in ddH_2_O. 0.6 μl of cell suspension was spotted onto slides and examined immediately. Cells were visualized and photographed through a Nikon Plan Apo VC 100×/1.40 oil objective lens by using a Nikon ECLIPSE 80i microscope (Nikon, Japan) equipped with a SPOT-RT KE monochrome charge-coupled device camera (Diagnostic Instruments, Sterling Heights, MI). Digital images were processed with Photoshop (Adobe system, Mountain View, CA).

For the temperature shift or knock sideways experiments, early-log phase cells (OD_660_ = ∼0.5) were incubated at 37°C for the indicated periods, plus or minus rapamycin for the indicated times. After incubation, cells were chilled on ice and washed by ice-cold ddH_2_O for once and then resuspended in appreciate amount of ice-cold ddH_2_O. 0.6 μl of cell suspension was spotted onto slides and examined immediately.

The staining of limiting membrane of the yeast vacuole was achieved with FM™ 4-64 Dye (N-(3-Triethylammoniumpropyl)-4-(6-(4-(Diethylamino) Phenyl) Hexatrienyl) Pyridinium Dibromide) (Invitrogen). Cells were grown to early-log phase in appropriate medium. Cells from 0.5ml of culture were resuspended in 50 μl of YEPD medium with 20 μM FM4-64. The cells were incubated for 20 minutes in a 30°C incubator or for 30 minutes in a 25°C incubator. Cells were then washed once with YEPD and resuspended in 1ml of YEPD. After incubating with vigorous shaking at 25°C for 90 to 120 min, cells were examined immediately.

For the <u>b</u>imolecular interaction fluorescent <u>c</u>omplementation (BiFC) assay, BY4741 cells are transformed with the indicated CEN plasmids and grown in double selection medium until mid-log phase (OD_660_ = ∼0.6). Cells were harvest by centrifugation and washed with ddH_2_O and resuspended in ddH_2_O. 0.6 μl of the cell suspension was spotted onto slides and examined immediately.

#### Confocal fluorescence microscopy

Yeast cells were grown in appropriate medium at 25°C to mid-log phase (OD_660_ = 0.6-0.8). Cells were harvest by centrifugation, resuspended in 3.7% paraformaldehyde (PFA) in phosphate buffered saline (PBS) and incubated for 20 minutes at room temperature. Following incubation, cells were washed once with PBS and resuspended in PBS. 0.6 μl of cell suspension was spotted onto Concanavalin A coated slides and examined using a Leica Sp8 Confocal Microscope. Cells were visualized and photographed through a Leica HCX PL APO 100×/1.40 oil objective lens using a Leica DMi8 CS Motorized inverted microscope equipped with HyD SP GaAsP detectors. Images were acquired with pixel sizes of 75nm. Due to low fluorescence signals from cells, final images were the result of the average of 8 individual images. Adobe Photoshop was used to adjust brightness and merge the images. *In vitro pull-down assays*

For *in vitro* protein pull down experiments yeast cells were grown to an OD_660_ of 1.0 and resuspended at 50 OD_660_ per mL in Buffer P (PBS and 0.1% Triton-X-100) supplemented with 1× EDTA-free complete protease inhibitor, 1 mM Pefabloc and 2mM DTT. Yeast cells were lysed using acid washed glass beads and a Vortex (). Following lysis homogenates were subjected to centrifugation at 13000×g for 10 min at 4°C. The resulting supernatant served as the yeast whole cell extracts for the pull-down experiments.

For mammalian pull down experiments, human 293T cells were lysed *in situ* by the addition of lysis buffer (20 mM HEPES pH 7.6, 175 mM NaCl, 0.25% NP-40, 10% glycerol, 1 mM EDTA, 1 mM DTT 1x protease inhibitor cocktail and 1 mM Pefabloc). Cell lysates were transferred to centrifugation tubes and incubated on ice for 30 min. After incubation, cell debris was removed by centrifugation at 16000×g for 10 min at 4°C. The resulting supernatant served as the mammalian whole cell extracts for the pull-down experiments.

Whole cell extracts from yeast or mammalian cells were incubated with purified recombinant yeast proteins prepared as described above. Lysates were added to a suspension of glutathione Sepharose 4B beads pre-equilibrated with Buffer A (25 mM Tris-HCl, pH 7.5, 400 mM KCl, 10% glycerol, 4% Triton X-100, 1xProtease inhibitor and 5mM DTT) and bound to bacterially expressed yeast proteins of interest at 4°C for 2 h with continuous gentle rotation. The beads were then washed 2 times with Buffer A and 2 times with Buffer C (25 mM Tris-HCl, pH 7.5, 150 mM KCl, 10% glycerol, 1% Triton X-100). GST-fusion proteins bound to glutathione Sepharose 4B beads were incubated with or without purified recombinant Sly1-His protein for 2 hours at 4°C. After 4 washes with buffer P, fusion proteins bound to beads were incubated with either purified yeast coatomer, yeast whole cell extract or mammalian whole cell extracts for 2 hours at 4°C. After incubation, the beads were washed four times with Buffer P, and bound proteins were identified following SDS– PAGE by immunoblotting. Recombinant proteins were visualized by Ponceau S staining of nitrocellulose membranes.

#### Assay for determining the steady-state level of individual proteins from yeast whole cell extracts

For detection of steady state protein levels from yeast cells, cells were grown at 25°C to mid-log phase (OD_660_ = ∼0.6), Lysates from cells grown at temperatures other than 25°C cultures were grown for 2 hours at the desired temperature. 10^7^ cells were harvest by centrifugation, resuspended in 200 μl of 0.1M of NaOH and incubated at room temperature for 5 minutes. Then cells were then collected by centrifugation, resuspended in 100 μL of SDS sample buffer containing 1× EDTA-free complete protease inhibitor, 1 mM Pefabloc and 5 mM DTT and incubated at 95°C for 5 minutes. Proteins were resolved by SDS-PAGE and detected by immunoblotting.

#### In vitro binding assays

Were performed essentially as described (Peng et al. 2002).

#### Native PAGE

Purified proteins prepared in standard buffer incubated at 4°C with gentle rotation overnight. After incubation, proteins were resolved by either SDS-PAGE or native PAGE and visualized by staining with Coomassie Briliant Blue.

#### Statistical analysis

Statistical analyses were carried out using Excel software (Microsoft Corporation). P-values were calculated using two-tail t-test of two samples assuming unequal variances.

## ACKNOWLEDGEMENTS

This work was supported by grants from the Hong Kong Research Council GRF 16101718 to DKB and by AoE/M-05/12-2. We thank members of the Banfield lab for feedback on the manuscript.

---

**Figure S1.**
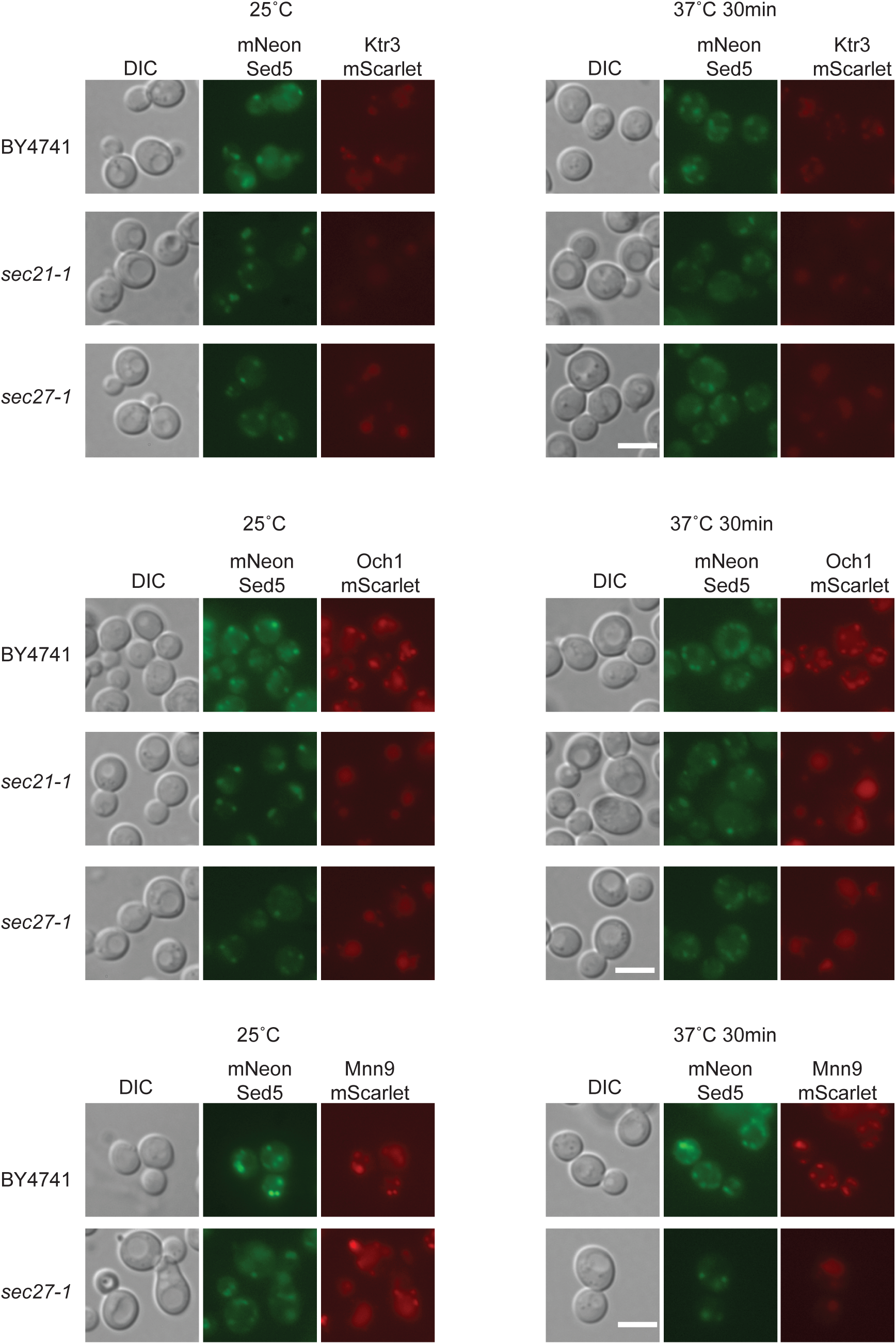
Golgi resident glycosyltransferases are mislocalized to the vacuole in COPI-coatomer temperature-sensitive mutants. BY4741 (wild-type), *sec21-1* and *sec27-1* cells expressing the indicated glycosyltransferases (see Table 1) were grown in YEPD, mounted on slides and photographed immediately. Scale bar 5 μm.

**Figure S2.**
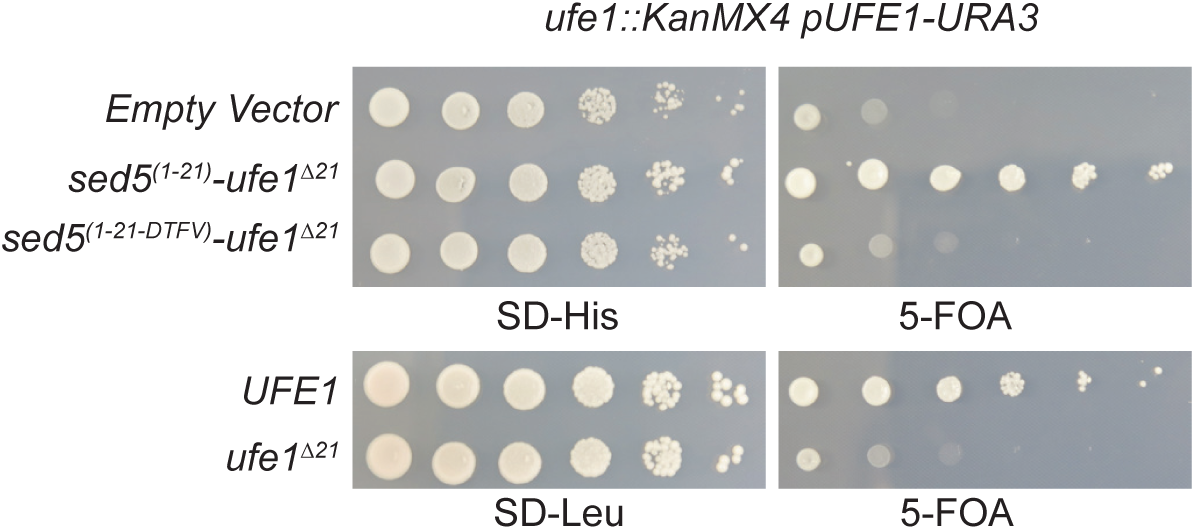
An N-terminal Sly1p-binding motif is essential for Ufe1p function. Cells containing a chromosomal deletion of *UFE1* and a copy of *UFE1* on a *URA3* containing plasmid (PRS316) were transformed with the indicated plasmids and 10-fold serial dilutions of transformants were spotted onto plates lacking either histidine (-His) or leucine (-Leu) and onto plates containing 5-FOA to counter-select for wildtype *UFE1* (pUFE1-URA3).

**Figure S3.**
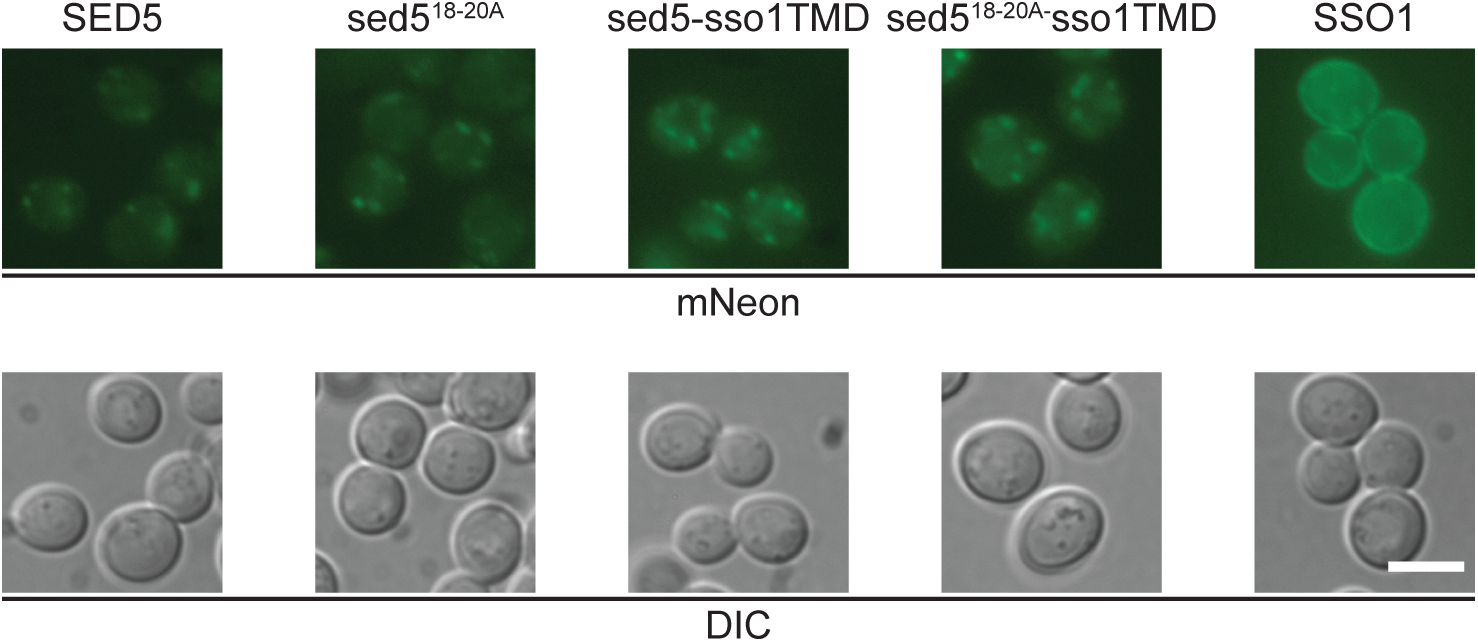
The Sed5-Sso1(TMD) chimeric protein lacking the COPI-coatomer tribasic motif is still localized to the Golgi. SEY6210 (wild-type) cells carrying chromosomal integrations of the indicated Sed5-Sso1 chimera at the *URA3* locus (see Table 1) were spotted into slides and photographed immediately. mNeon-Sso1p localizes to the limiting membrane of cell whereas all other constructs resemble mNeon-Sed5p and localize to puncta. Scale bar 5 μm.

**Figure S4.**
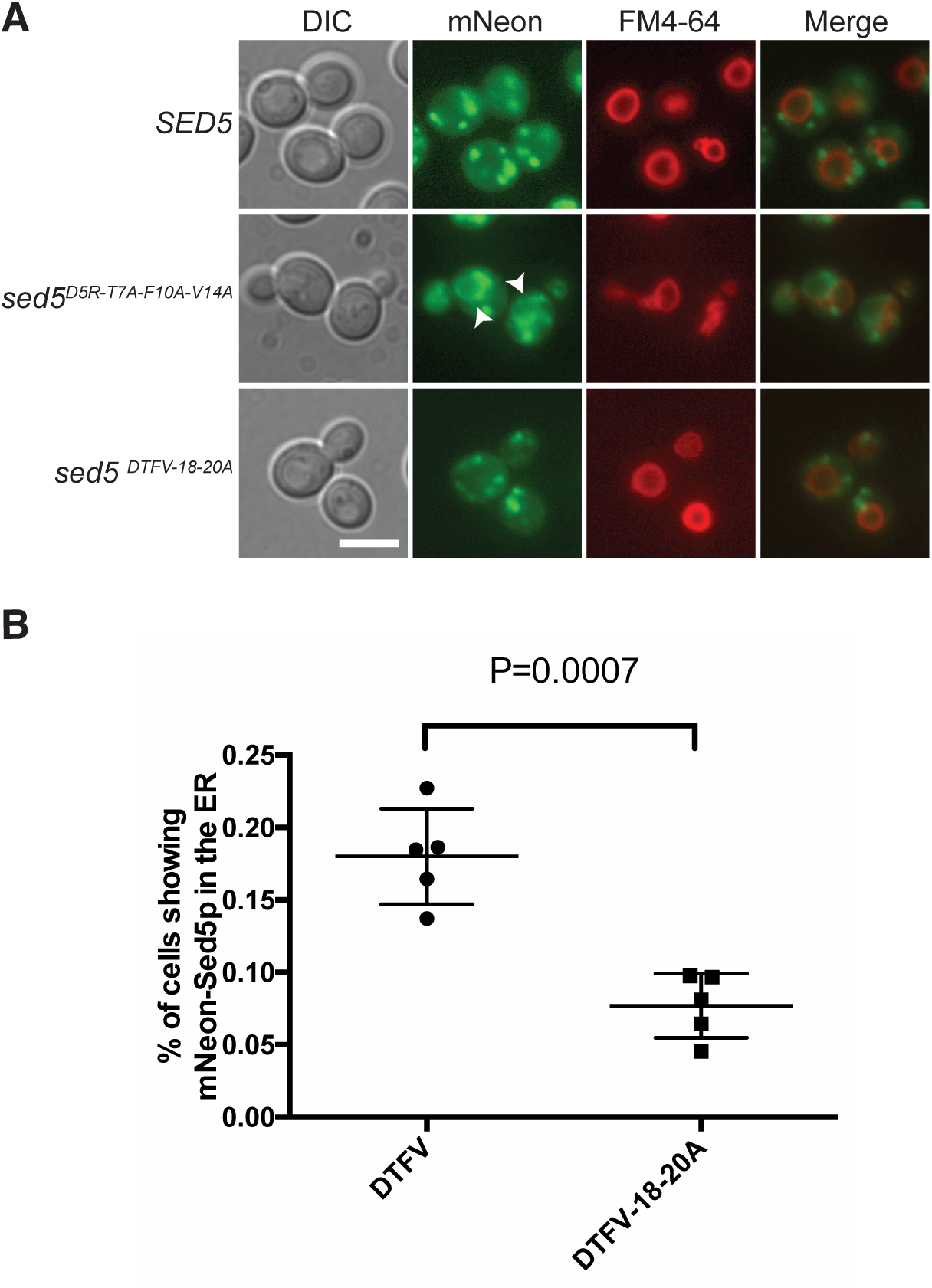
The Golgi localization of Sed5p is independent of Sly1p and COPI-coatomer binding. mNeon *SED5*, mNeon *sed5* DTFV or mNeon *sed5* DTFV 18-20A were integrated at the *URA3* locus in SEY6210 cells (see Table 1). mNeon sed5p DTFV localizes to both puncta and the ER, all other constructs localize primarily to punta. Quantification of the percentage of mNeon *sed5* DTFV and mNeon *sed5* DTFV 18-20A cells showing ER staining is depicted below. Mean ± SD *P = 0.0007 calculated by unpaired Student’s t-test assuming Unequal Variances; N = 5 isolates, n>500. Scale bar 5 μm.

**Figure S5.**
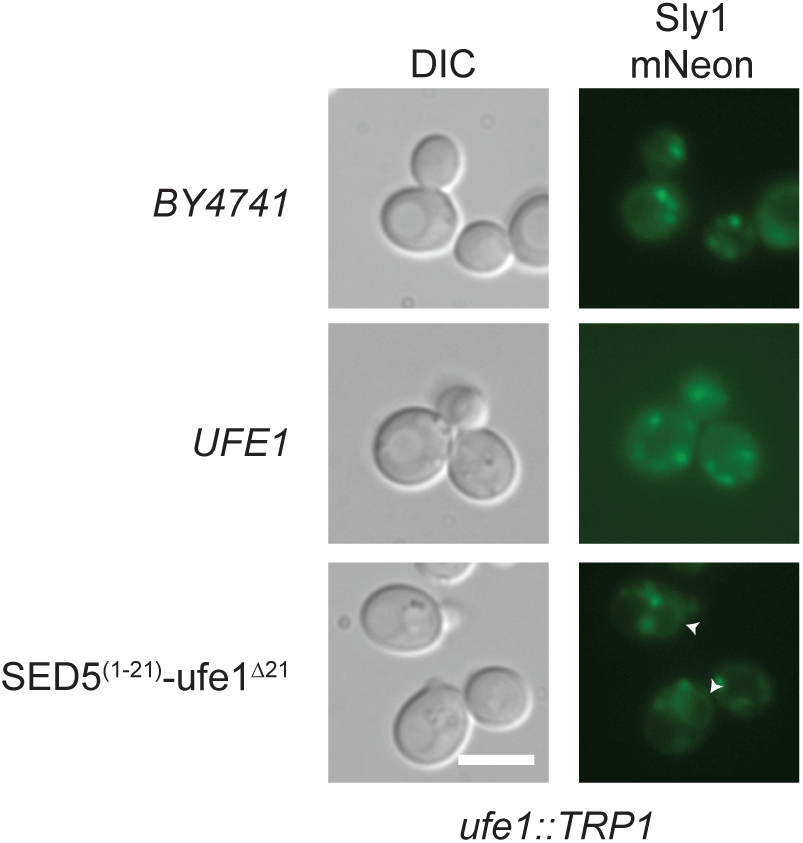
In cells expressing a Sed5p-Ufe1p chimera Sly1p is localized to Golgi and ER. Widltype cells (BY4741) and cells lacking endogenous *UFE1* (*ufe1::TRP1*) were transformed with the indicated plasmids and wildtpe *UFE1* was removed by counter-selection on plates containing 5-FOA. Cells were cultured post 5-FOA treatment and grown to early log phase in liquid media thereafter cells were mounted onto slides and photographed immediately. Arrowheads denote the nuclear envelop / ER. Scale bar 5μm.

**Figure S6.**
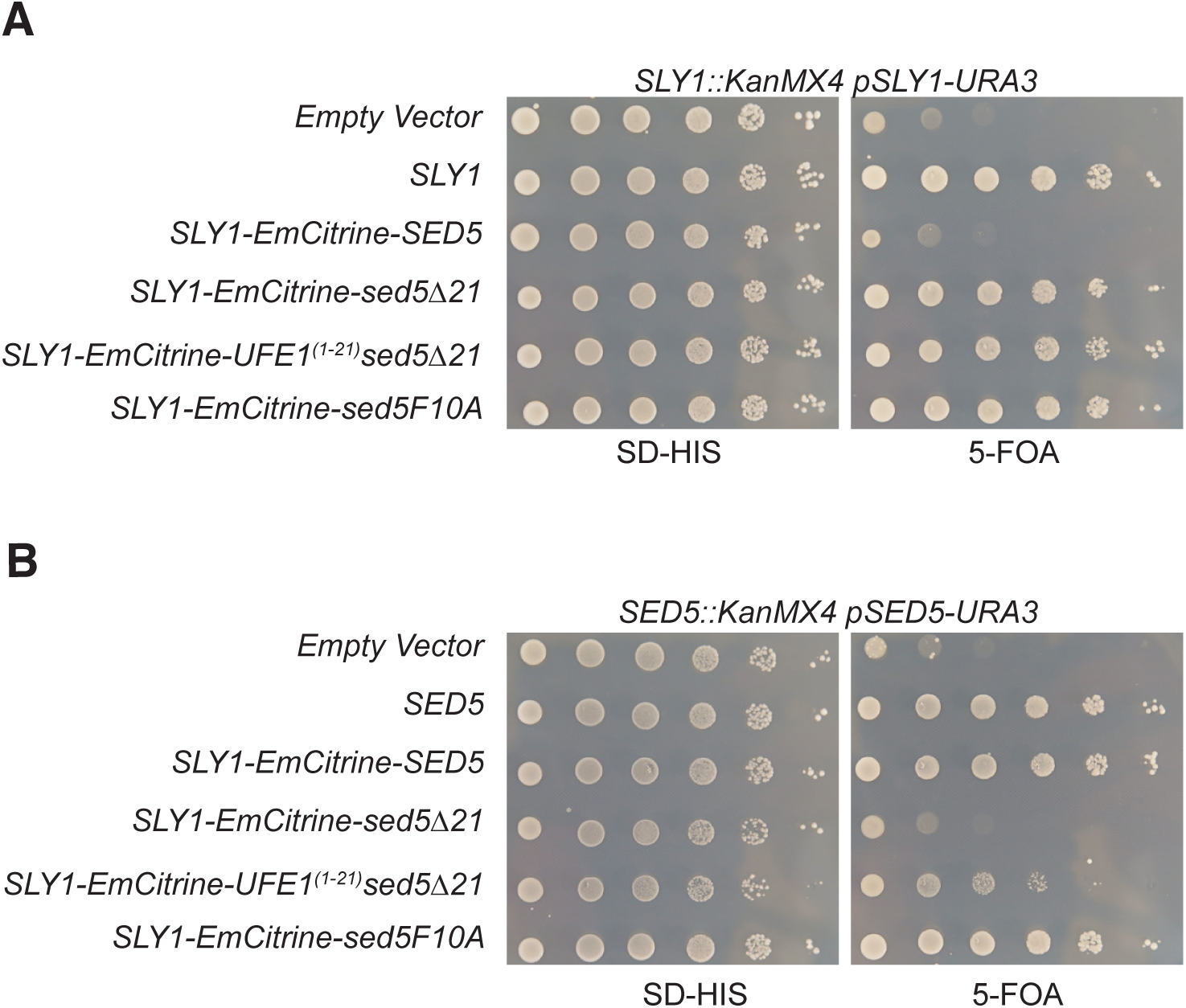
Functional requirements of Sly1p-emCitrine-Sed5p in sly1Δ and sed5Δ cells. (A) A weakened interaction between Sly1p and Sed5p is necessary for a covalent protein (Sly1p-emCitrine-Sed5p) to support the growth of *sly1Δ* cells (B) The first 21 amino acids of Sed5p are essential for the function of Sly1p-emCitrine-Sed5p in *sed5Δ* cells. The indicated balanced single deletion strains were transformed with the various constructs listed on the left hand side of each panel. Transformants were plated as 10-fold serial dilutions and incubated at 25°C for 2 days prior to being photographed.

## REFERENCES

Aalto MK, Ronne H & Keränen S (1993) Yeast syntaxins Sso1p and Sso2p belong to a family of related membrane proteins that function in vesicular transport. EMBO J. 12: 4095–104

Arakel EC & Schwappach B (2018) Formation of COPI-coated vesicles at a glance. J. Cell Sci. 131: jcs209890

Aridor M & Traub LM (2002) Cargo selection in vesicular transport: The making and breaking of a coat. Traffic 3: 537–546

Baker RW, Jeffrey PD, Zick M, Phillips BP, Wickner WT & Hughson FM (2015) A direct role for the Sec1/Munc18-family protein Vps33 as a template for SNARE assembly. Science (80-.). 349: 1111–1114

Ballensiefen W, Ossipov D & Schmitt HD (1998) Recycling of the yeast v-SNARE Sec22p involves COPI-proteins and the ER transmembrane proteins Ufe1p and Sec20p. J. Cell Sci. 111: 1507 LP–1520

Banfield DK (2011) Mechanisms of protein retention in the Golgi. Cold Spring Harb. Perspect. Biol. 3: 1–14

Banfield DK, Lewis MJ & Pelham HRB (1995) A SNARE-like protein required for traffic through the Golgi complex. Nature 375: 806–809

Banfield DK, Lewis MJ, Rabouille C, Warren G & Pelham HRB (1994) Localization of Sed5, a putative vesicle targeting molecule, to the cis-Golgi network involves both its transmembrane and cytoplasmic domains. J. Cell Biol. 127: 357–371

Bracher A & Weissenhorn W (2002) Structural basis for the Golgi membrane recruitment of Sly1p by Sed5p. EMBO J. 21: 6114–6124

Braun S & Jentsch S (2007) SM-protein-controlled ER-associated degradation discriminates between different SNAREs. EMBO Rep. 8: 1176–1182

Demircioglu FE, Burkhardt P & Fasshauer D (2014) The SM protein Sly1 accelerates assembly of the ER-Golgi SNARE complex. Proc. Natl. Acad. Sci. 111: 13828–13833

Dulubova I, Sugita S, Hill S, Hosaka M, Fernandez I, Südhof TC, Rizo J, Advani R, Bae H, Bock J, Chao D, Doung Y, Prekeris R, Yoo J, Scheller R, Banerjee A, Kowalchyk J, DasGupta B, Martin T, Bennett M, et al (1999) A conformational switch in syntaxin during exocytosis: role of munc18. EMBO J. 18: 4372–4382

Fasshauer D, Sutton RB, Brunger AT & Jahn R (1998) Conserved structural features of the synaptic fusion complex: SNARE proteins reclassified as Q- and R-SNAREs. Proc. Natl. Acad. Sci. 95: 15781–15786

Gerst JE, Rodgers L, Riggs M & Wigler M (1992) SNC1, a yeast homolog of the synaptic vesicle-associated membrane protein/synaptobrevin gene family: Genetic interactions with the RAS and CAP genes. Proc. Natl. Acad. Sci. U. S. A. 89: 4338–4342

Glick BS, Elston T & Oster G (1997) A cisternal maturation mechanism can explain the asymmetry of the Golgi stack. FEBS Lett. 414: 177–181

Grabowski R & Gallwitz D (1997) High-affinity binding of the yeast cis-Golgi t-SNARE; Sed5p, to wild-type and mutant Sly1p, a modulator of transport vesicle docking. FEBS Lett. 411: 169–172

Graf CT, Riedel D, Schmitt HD & Jahn R (2005) Identification of functionally interacting SNAREs by using complementary substitutions in the conserved ‘0’ layer. Mol Biol Cell 16: 2263–2274

Gurunathan S, Chapman-Shimshoni D, Trajkovic S & Gerst JE (2000) Yeast exocytic v-SNAREs confer endocytosis. Mol. Biol. Cell 11: 3629–3643

Hardwick KG & Pelham HRB (1992) SED5 encodes a 39-kD integral membrane protein required for vesicular transport between the ER and the Golgi complex. J. Cell Biol. 119: 513–521

Hui N, Nakamura N, Sönnichsen B, Shima DT, Nilsson T & Warren G (1997) An isoform of the Golgi t-SNARE, syntaxin 5, with an endoplasmic reticulum retrieval signal. Mol. Biol. Cell 8: 1777–87

Jäntti J, Aalto MK, Oyen M, Sundqvist L, Keränen S & Ronne H (2002) Characterization of temperature-sensitive mutations in the yeast syntaxin 1 homologues Sso1p and Sso2p, and evidence of a distinct function for Sso1p in sporulation. J. Cell Sci. 115: 409–20

Kerppola TK (2006) Design and implementation of bimolecular fluorescence complementation (BiFC) assays for the visualization of protein interactions in living cells. Nat. Protoc. 1: 1278–86

Koike S & Jahn R (2019) SNAREs define targeting specificity of trafficking vesicles by combinatorial interaction with tethering factors. Nat. Commun.

Kurokawa K, Okamoto M & Nakano A (2014) Contact of cis-Golgi with ER exit sites executes cargo capture and delivery from the ER. Nat. Commun. 5: 3653

Lee MCS, Miller EA, Goldberg J, Orci L & Schekman R (2004) BI-DIRECTIONAL PROTEIN TRANSPORT BETWEEN THE ER AND GOLGI. Annu. Rev. Cell Dev. Biol. 20: 87–123

Lewis MJ, Rayner JC & Pelham HRB (1997) A novel SNARE complex implicated in vesicle fusion with the endoplasmic reticulum. EMBO J. 16: 3017–3024

Li Y, Gallwitz D & Peng R (2005) Structure-based functional analysis reveals a role for the SM protein Sly1p in retrograde transport to the endoplasmic reticulum. Mol. Biol. Cell 16: 3951–3962

Liu L, Doray B & Kornfeld S (2018) Recycling of Golgi glycosyltransferases requires direct binding to coatomer. Proc. Natl. Acad. Sci. U. S. A. 115: 8984–8989

Lobingier BT, Nickerson DP, Lo SY & Merz AJ (2014) SM proteins Sly1 and Vps33 co-assemble with Sec17 and SNARE complexes to oppose SNARE disassembly by Sec18. Elife 2014:

Losev E, Reinke CA, Jellen J, Strongin DE, Bevis BJ & Glick BS (2006) Golgi maturation visualized in living yeast. Nature 441: 1002–1006

Matsuura-Tokita K, Takeuchi M, Ichihara A, Mikuriya K & Nakano A (2006) Live imaging of yeast Golgi cisternal maturation. Nature 441: 1007–1010

McNew JA, Coe JG s, Søgaard M, Zemelman B V, Wimmer C, Hong W & Söllner TH (1998) Gos1p, a Saccharomyces cerevisiae SNARE protein involved in Golgi transport. FEBS Lett. 435: 89–95

McNew JA, Søgaard M, Lampen NM, Machida S, Ye RR, Lacomis L, Tempst P, Rothman JE & Söllner TH (1997) Ykt6p, a prenylated SNARE essential for endoplasmic reticulum-golgi transport. J. Biol. Chem. 272: 17776–17783

Miller EA, Liu Y, Barlowe C & Schekman R (2005) ER-Golgi transport defects are associated with mutations in the Sed5p-binding domain of the COPII coat subunit, Sec24p. Mol. Biol. Cell 16: 3719–3726

Miyazaki K, Wakana Y, Noda C, Arasaki K, Furuno A & Tagaya M (2012) Contribution of the long form of syntaxin 5 to the organization of the endoplasmic reticulum. J. Cell Sci. 125: 5658–5666

Mossessova E, Bickford LC & Goldberg J (2003) SNARE selectivity of the COPII coat. Cell 114: 483–495

Muhlrad D, Hunter R & Parker R (1992) A rapid method for localized mutagenesis of yeast genes. Yeast 8: 79–82

Munro S (1995) An investigation of the role of transmembrane domains in Golgi protein retention. EMBO J. 14: 4695–4704

Nakanishi H, Morishita M, Schwartz CL, Coluccio A, Engebrecht JA & Neiman AM (2006) Phospholipase D and the SNARE Sso1p are necessary for vesicle fusion during sporulation in yeast. J. Cell Sci. 119: 1406–1415

Opat AS, Van Vliet C & Gleeson PA (2001) Trafficking and localisation of resident Golgi glycosylation enzymes. Biochimie 83: 763–773

Ossipov D, Schröder-Köhne S & Schmitt HD (1999) Yeast ER-Golgi v-SNAREs Bos1p and Bet1p differ in steady-state localization and targeting. J. Cell Sci.

Papanikou E, Day KJ, Austin J & Glick BS (2015) COPI selectively drives maturation of the early Golgi. Elife 4:

Parlati F, McNew JA, Fukuda R, Miller R, Söllner TH & Rothman JE (2000) Topological restriction of SNARE-dependent membrane fusion. Nature 407: 194–198

Parlati F, Varlamov O, Paz K, McNew JA, Hurtado D, Söllner TH & Rothman JE (2002) Distinct SNARE complexes mediating membrane fusion in Golgi transport based on combinatorial specificity. Proc. Natl. Acad. Sci. U. S. A. 99: 5424–5429

Peng R & Gallwitz D (2004) Multiple SNARE interactions of an SM protein: Sed5p/Sly1p binding is dispensable for transport. EMBO J. 23: 3939–3949

Pryor PR, Jackson L, Gray SR, Edeling MA, Thompson A, Sanderson CM, Evans PR, Owen DJ & Luzio JP (2008) Molecular Basis for the Sorting of the SNARE VAMP7 into Endocytic Clathrin-Coated Vesicles by the ArfGAP Hrb. Cell 134: 817–827

Rayner JC & Pelham HRB (1997) Transmembrane domain-dependent sorting of proteins to the ER and plasma membrane in yeast. EMBO J. 16: 1832–1841

Reggiori F, Black MW & Pelham HRB (2000) Polar transmembrane domains target proteins to the interior of the yeast vacuole. Mol. Biol. Cell 11: 3737–3749

Rothman JE & Warren G (1994) Implications of the SNARE hypothesis for intracellular membrane topology and dynamics. Curr. Biol. 4: 220–33

Sacher M, Stone S & Ferro-Novick S (1997) The synaptobrevin-related domains of Bos1p and Sec22p bind to the syntaxin-like region of Sed5p. J. Biol. Chem. 272: 17134–17138

Sato K, Sato M & Nakano A (2001) Rer1p, a retrieval receptor for endoplasmic reticulum membrane proteins, is dynamically localized to the Golgi apparatus by coatomer. J. Cell Biol. 152: 935–944

Schmitz KR, Liu J, Li S, Setty TG, Wood CS, Burd CG & Ferguson KM (2008) Golgi Localization of Glycosyltransferases Requires a Vps74p Oligomer. Dev. Cell 14: 523–534

Schröter S, Beckmann S & Schmitt HD (2016) ER arrival sites for COPI vesicles localize to hotspots of membrane trafficking. EMBO J. 35: 1935–1955

Sharpe HJ, Stevens TJ & Munro S (2010) A Comprehensive Comparison of Transmembrane Domains Reveals Organelle-Specific Properties. Cell 142: 158–169

Søgaard M, Tani K, Ye RR, Geromanos S, Tempst P, Kirchhausen T, Rothman JE & Söllner T (1994) A rab protein is required for the assembly of SNARE complexes in the docking of transport vesicles. Cell 78: 937–948

Söllner T, Whiteheart SW, Brunner M, Erdjument-Bromage H, Geromanos S, Tempst P & Rothman JE (1993) SNAP receptors implicated in vesicle targeting and fusion. Nature 362: 318–324

Tochio H, Tsui MMK, Banfield DK & Zhang M (2001) An autoinhibitory mechanism for nonsyntaxin SNARE proteins revealed by the structure of Ykt6p. Science (80-.). 293: 698–702

Tsui MMK, Tai WCS & Banfield DK (2001) Selective Formation of Sed5p-containing SNARE Complexes Is Mediated by Combinatorial Binding Interactions. Mol. Biol. Cell 12: 521–538

Tu L, Chen L & Banfield DK (2012) A Conserved N-terminal Arginine-Motif in GOLPH3-Family Proteins Mediates Binding to Coatomer. Traffic 13: 1496–1507

Tu L, Tai WCS, Chen L & Banfield DK (2008) Signal-mediated dynamic retention of glycosyltransferases in the Golgi. Science 321: 404–7

Valdez-Taubas J & Pelham HRB (2003) Slow diffusion of proteins in the yeast plasma membrane allows polarity to be maintained by endocytic cycling. Curr. Biol. 13: 1636–1640

Villeneuve J, Duran J, Scarpa M, Bassaganyas L, Galen J Van & Malhotra V (2017) Golgi enzymes do not cycle through the endoplasmic reticulum during protein secretion or mitosis. Mol. Biol. Cell 28: 141–151

Weber T, Zemelman B V, McNew JA, Westermann B, Gmachl M, Parlati F, Söllner TH & Rothman JE (1998) SNAREpins: Minimal Machinery for Membrane Fusion. Cell 92: 759–772

Weinberger A, Kamena F, Kama R, Spang A & Gerst JE (2005) Control of Golgi Morphology and Function by Sed5 t-SNARE Phosphorylation. Mol. Biol. Cell 16: 4918–4930

Wen W, Chen L, Wu H, Sun X, Zhang M & Banfield DK (2006) Identification of the yeast R-SNARE Nyv1p as a novel longin domain-containing protein. Mol. Biol. Cell 17: 4282–4299

Wood CS, Schmitz KR, Bessman NJ, Setty TG, Ferguson KM & Burd CG (2009) PtdIns4P recognition by Vps74/GOLPH3 links PtdIns 4-kinase signaling to retrograde Golgi trafficking. J. Cell Biol. 187: 967–975

Wooding S & Pelham HRB (1998) The dynamics of Golgi protein traffic visualized in living yeast cells. Mol. Biol. Cell 9: 2667–2680

Yamaguchi T, Dulubova I, Min SW, Chen X, Rizo J & Südhof TC (2002) Sly1 binds to Golgi and ER syntaxins via a conserved N-terminal peptide motif. Dev. Cell 2: 295–305

Zink S, Wenzel D, Wurm CA & Schmitt HD (2009) A Link between ER Tethering and COP-I Vesicle Uncoating. Dev. Cell 17: 403–416

